# JASMINE: A powerful representation learning method for enhanced analysis of incomplete multi-omics data

**DOI:** 10.1101/2025.06.16.659949

**Authors:** Jenna L. Ballard, Zongyu Dai, Li Shen, Qi Long, Alzheimer’s Disease Neuroimaging Initiative, Alzheimer’s Disease Metabolomics Consortium

**Affiliations:** Graduate Group in Genomics and Computational Biology, Perelman School of Medicine, University of Pennsylvania, 3700 Hamilton Walk, Philadelphia, 19104, PA, USA; Graduate Group in Applied Mathematics and Computational Science, University of Pennsylvania, 209 S. 33rd Street, Philadelphia, 19104, PA, USA; Department of Biostatistics, Epidemiology and Informatics, Perelman School of Medicine, University of Pennsylvania, 423 Guardian Drive, Philadelphia, 19104, PA, USA

**Keywords:** representation learning, variational autoencoder, multimodal integration, missing data

## Abstract

Integrative analysis of multi-omics data provides a more comprehensive and nuanced view of a subject’s biological state. However, high-dimensionality and ubiquitous modality missingness present significant analytical challenges. Existing methods for incomplete multi-omics data are scarce, do not fully leverage both modality-specific and shared information, and produce task-biased representations. We propose JASMINE, a self-supervised representation learning method for incomplete multi-omics data that preserves both modality-specific and joint information and enhances sample similarity structure. JASMINE produces embeddings that achieve superior performance across multiple tasks for two different incomplete multi-omics datasets while requiring only a single round of training per dataset.

## 1 Background

With advances in data collection technologies, storage and computing resources, vast amounts of -omics data, including genetic variation (genomics), gene regulation (epigenomics), gene expression (transcriptomics), protein abundance (proteomics), and metabolite abundance (metabolomics), are becoming available. Each of these modalities sheds light on various biological processes. Multiple modalities are often measured for the same subject, providing more comprehensive information which may aid downstream analyses and improve performance on predictive tasks. One application of interest is to disease pathophysiology, which is complex and multifaceted, resulting from the interactions between processes across many layers, including the molecular, cellular, tissue, organ, and patient levels. Combining the information offered by diverse -omics data modalities may better capture these processes and their interactions which underlie particular phenotypes. This can enhance diagnosis [1], disease subtyping [2], and discovery of disease-relevant biological pathways [3, 4].

Common obstacles to fully leveraging multi-omics data include high dimensionality and incompleteness: some modalities can comprise tens of thousands of features, and most datasets contain a considerable proportion of samples with missing modalities. The ability to overcome these challenges necessitates strategies to produce clean, compact representations for each sample that are amenable to downstream analyses and prediction tasks.

Many strategies have been devised to integrate multi-omics data for downstream tasks. These can be broadly categorized as early, intermediate, or late integration. Early-integration combines the modalities as a preprocessing step, such as by concatenating the features across data types and using this as input to the model [5–7]. Late integration applies separate models to each data modality and then aggregates the outputs into a final multi-modal prediction [8–10]. These approaches, which handle individual modalities by either simply concatenating their features before inputting them to a model or by applying separate models, may better preserve modality-specific information [5], but they are limited in their ability to fully learn and exploit inter-modality interactions. On the other hand, intermediate integration strategies recognize the modalities as distinct entities at the input level, and then fuse their information while learning inter-modality relationships [5, 11–15].

Another important consideration for multi-omics integration is the information preserved by the representations. Two principles of interest are consensus, which holds that model errors are upper-bounded by the level of disagreement between modalities, and complementarity, which holds that each modality contains critical unique information [5]. Early- and late-integration approaches tend to better preserve modality-specific information, satisfying the complementarity principle. Intermediate integration-based methods can satisfy both complementarity and consensus [16–18] by including modality-specific and shared components, respectively. Some strategies [16, 18] further ensure that the two components contain unique information by imposing orthogonality constraints between them.

Despite the availability of methods for integrating multi-omics data, most leave the issue of missing modalities unaddressed. A common way to handle this is to exclude samples with incomplete data. However, this reduces the sample size which may lead to reduced power in subsequent analysis; and the remaining subset of subjects may not be representative of the original dataset which may lead to biased results. Other techniques impute the missing data as a preprocessing step, but this may bias the imputed values to be similar, which can negatively impact downstream analyses [19]. Some recent multi-modal integration methods handle incomplete data by defining a joint latent space that can be derived from subsets of modalities [17, 20]. Such techniques include product-of-experts (PoE) and mixture-of-experts (MoE), which define the joint latent space as the product [13] and weighted average [21], respectively, of the modality-specific latent distributions. When modalities are missing, they are excluded from the calculation of the joint embedding distribution. Other methods can handle missing data by generating missing modalities from those that are available via cross-encoders [12] or enforcing similarity among the embeddings derived from all modalities [22]. A more recent method, IntegrAO [15], learns omics-specific patient graphs before fusing them to generate low dimensional embeddings in a unified space while using cross-patient similarities to help infer missing information. However, this graph-based method is memory intensive, becoming more prohibitive for both training and inference as sample size increases.

Other limitations of prior methods is that they are often supervised [20, 23–26], resulting in representations that are biased towards a specific task, and requiring retraining for different tasks. The goal of our method is to generate informative, compact representations of multi-omics data that attain superior performance on a variety of downstream tasks. To do this, we use a self-supervised approach based on reconstruction loss. Self-supervised learning is similar to unsupervised learning in that it learns inherent patterns in the data without utilizing external labels. However, a crucial distinction between the two is that only self-supervised learning uses a ground truth target for optimization. In our case, the target is the original input which the model aims to be able to reconstruct by minimizing reconstruction loss. Reconstruction-based self-supervision trains the method to distill the high-dimensional input data into low-dimensional embeddings that retain sufficient information to recover the original data. Because these embeddings have many fewer features than the original data, minimizing reconstruction loss ensures that the retained features contain only the most essential information in the data. As a result, any irrelevant noise present in the high-dimensional input is reduced. Therefore, this approach has important functions both in dimensionality reduction and in data denoising, while extracting patterns intrinsic to the data. Supervised approaches, on the other hand, focus on learning the relationship between the input data and an external label, which may cause the model to ignore important information that is not relevant to the specific prediction task for which it is being trained. By training without a particular prediction task objective, our approach produces task-agnostic representations that excel at multiple supervised tasks while also retaining flexibility for unsupervised tasks such as clustering.

To accomplish these goals, we developed JASMINE (**J**oint **A**nd modality-**S**pecific **M**ultimodal representation learning handling **IN**compl**E**te data), a self-supervised representation learning method that generates compact, task-agnostic embeddings that integrate multi-omics data while handling arbitrarily missing modalities. The main contributions of this work are the following:

1. Specialized encoders and a PoE formulation are designed to produce modality-specific and joint components that preserve both complementary and shared information, respectively, among the modalities. Orthogonality constraints are used to minimize redundancy among these components.
2. Two types of contrastive learning (CL), at the modality level and sample level, enhance the discriminative structure of the data while reducing irrelevant noise. Modality-level CL enforces consistency among the modalities for a given sample, and sample-level CL utilizes a deep clustering-based approach to ensure that similarity relationships between samples in the original feature space are preserved in the embedding space.
3. Samples with arbitrarily missing modalities are handled using both cross-encoders for modality-specific components and PoE for the joint component of the representation.
4. Self-supervised approach produces one set of representations that can be used for multiple different downstream tasks after a single round of training per dataset.
5. Simulation experiments demonstrated JASMINE’s suitability compared to baseline methods for cases when modalities are missing at higher rates, especially as the number of modalities increases and sample size is limited, and when missingness is unbalanced among ground truth classes.
6. Representations achieved best or close to the best performance on multiple prediction tasks across two different incomplete multi-omics datasets.

## 2 Results

### 2.1 JASMINE Framework

We present JASMINE, a variational autoencoder-based framework to learn representations of multi-omics data while handling missing modalities. Training is done in a self-supervised manner, resulting in representations that can be used for a variety of downstream tasks. JASMINE’s architecture and objective function are constructed to produce representations consisting of distinct components containing modality-specific and shared information. By incorporating self- and cross-encoders (see Section 5.2), the method is designed to preserve modality-specific information while learning pairwise cross-modality relationships. Reconstruction loss ensures that each resulting modality-specific component retains sufficient information to recover its corresponding input modality. To capture information shared among all modalities, the method additionally learns a joint representation using PoE (see Section 5.2.1), which defines the joint latent space as the product of the modality-specific distributions. Recon-struction loss for the joint component encourages it to preserve the information that is needed to reconstruct all modalities and thus is shared among them. The final representation is formed from concatenating the modality-specific and joint components (see Section 5.2.2). A schematic of JASMINE is presented in Figure 1. Our proposed approach further uses orthogonality constraints to minimize the redundancy of information between the modality-specific and shared components (see Section 5.2.3). It also utilizes modality-level contrastive learning (MCL) and sample-level contrastive learning (SCL) to encourage consistency among embeddings of different modalities from the same sample as well as preserve the similarity structure between samples in the embedding space (see Section 5.2.4). The final objective function comprises loss terms for the encoders, contrastive learning, and orthogonality constraints (see Section 5.2.5). Thus, our approach aims to generate informative representations that preserve both modality-specific and shared information while enhancing the similarity structure among samples. We evaluate the efficacy of the representations generated by our model on classification, clustering and regression tasks in simulated datasets and two different real datasets.

**Fig. 1.**
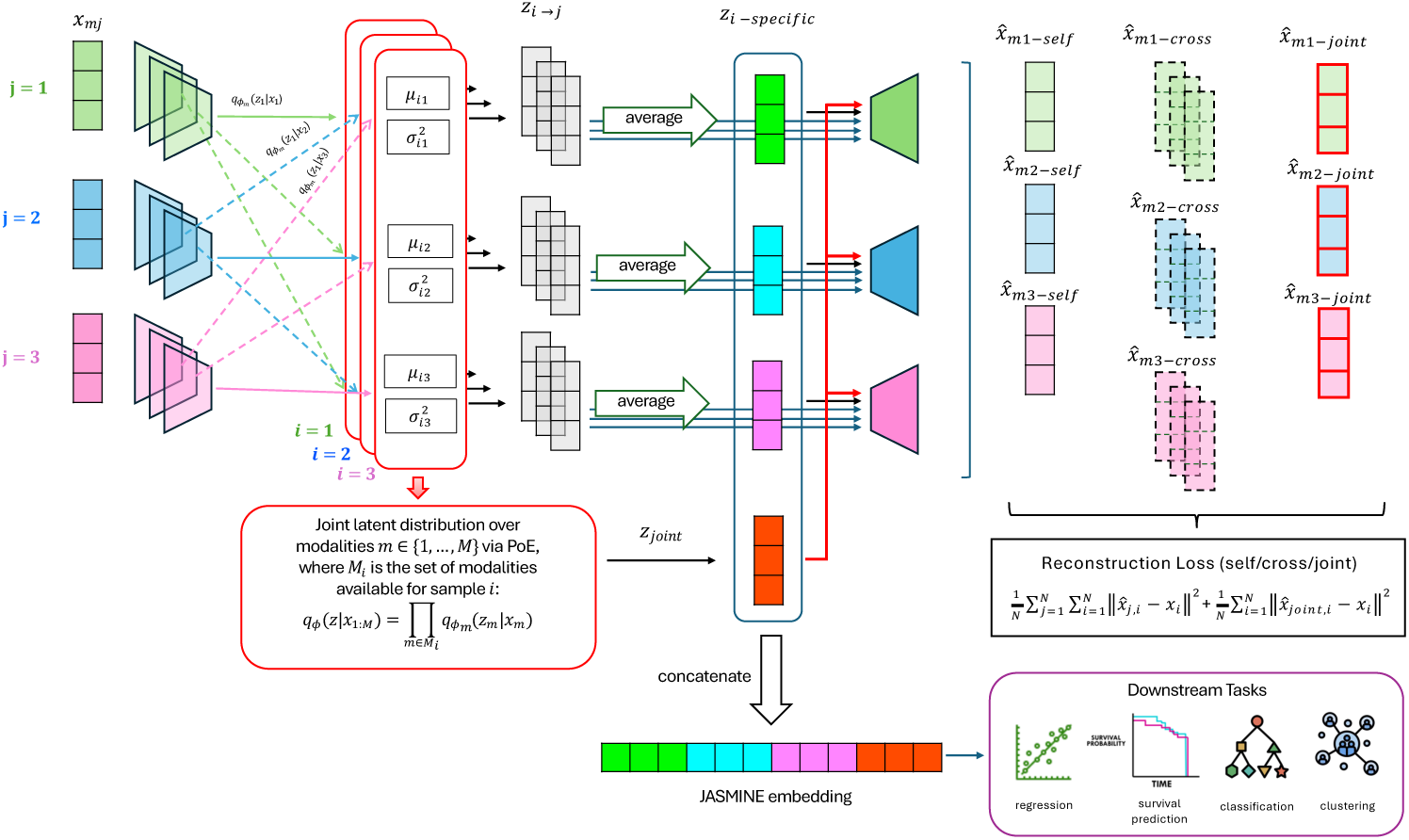
JASMINE Method Schematic. For data with *m* modalities, the model consists of *m*^2^ self- and cross-encoders which generate modality-specific representations. To get a single modality-specific representation for a given modality, the representations generated for that modality are averaged. A joint latent space is derived using PoE, from which a joint embedding is generated. Modality-specific and joint representations are concatenated to produce the final JASMINE embedding. These embeddings can be used as input for any desired downstream task (e.g. regression, classification, clustering). JASMINE is learned in a self-supervised fashion based on reconstruction loss. There is one decoder per modality for a total of *m* decoders, which are used to reconstruct each modality from its corresponding latent representation.

### 2.2 Experimental Setup

Our experimental pipeline consists of two main steps: (1) generate embeddings using an unsupervised representation learning method and (2) input the embeddings to a prediction model of choice for a given inference problem. In this way, the learned embeddings can be applied to various downstream tasks. More details on the data processing for this pipeline are given in Section 5.6. Here, we apply the embeddings to three types of prediction problems: (1) survival prediction using a Coxnet regression model; (2) classification using an XGBoost classification model (XGBClassifier); and (3) regression using an XGBoost regression model (XGBRegressor).

We compare JASMINE to multiple baseline embedding frameworks. The baselines include the original data, deep canonical correlation analysis (DCCA) [27], generalized canonical correlation analysis (GCCA) [28], multi-modal variational autoencoder (MVAE) [13], cross-linked unified embedding (CLUE) [12], and IntegrAO [15]. Details about these baseline methods can be found in Section 5.3. A summary of the key features of these methods can be found in Table 1. We evaluate and compare these methods based on various performance metrics. For survival prediction, we use the concordance index for right-censored data based on inverse probability of censoring weights (C-index IPCW) and cumulative dynamic area under the receiver operating characteristic curve (CD AUC). For classification, we use balanced accuracy, area under the receiver operating characteristic curve (AUROC) and average precision score (AP). In the multiclass setting, the macro-average across classes is utilized. For clustering, we assess differential survival via the log-rank test over various numbers of clusters and for five different cancer types. For regression, we used root mean squared error (RMSE), the coefficient of determination (*R*^2^-score), and the Pearson correlation between the predicted and true values (r-pred). More details about these metrics can be found in Section 5.4.

**Table 1.**
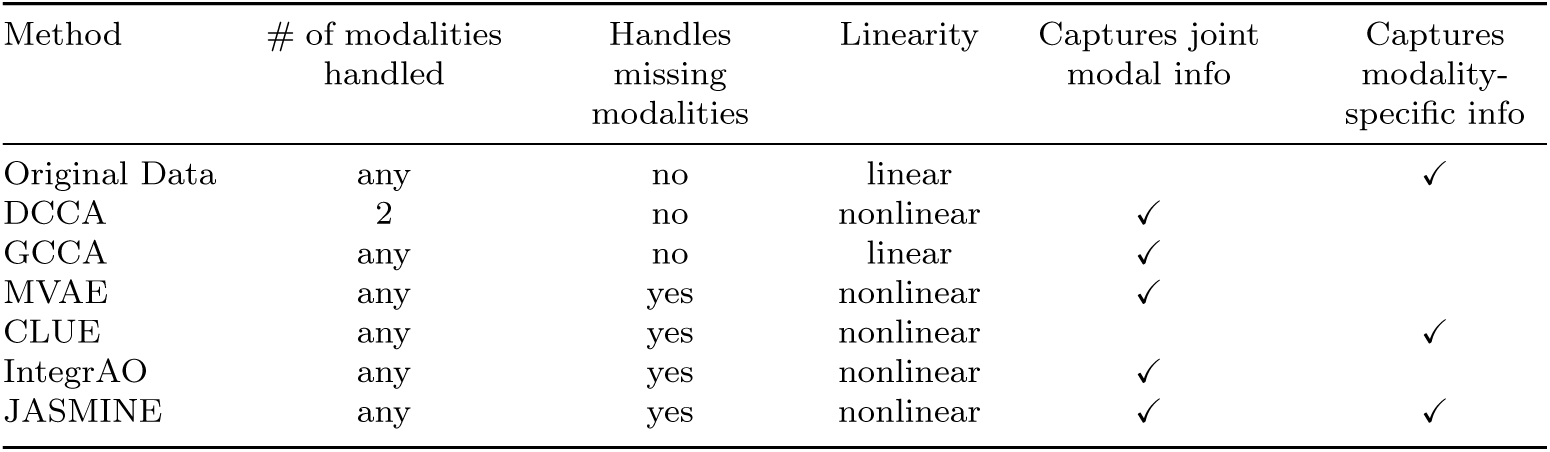
Proposed and Baseline Method Summary.

| Method | # of modalities handled | Handles missing modalities | Linearity | Captures joint modal info | Captures modality-specific info |
| --- | --- | --- | --- | --- | --- |
| Original Data | any | no | linear |  | ✓ |
| DCCA | 2 | no | nonlinear | ✓ |  |
| GCCA | any | no | linear | ✓ |  |
| MVAE | any | yes | nonlinear | ✓ |  |
| CLUE | any | yes | nonlinear |  | ✓ |
| IntegrAO | any | yes | nonlinear | ✓ |  |
| JASMINE | any | yes | nonlinear | ✓ | ✓ |

### 2.3 Evaluation on simulated data

To test JASMINE under a variety of controlled data settings, we evaluated and compared the performance of JASMINE and the baselines on simulated data. This included varying the level of modality missingness, evaluating the performance when missingness rates depended on the true label, and varying the number of modalities under small and large sample sizes. The synthetic data were simulated to comprise three distinct classes, each with 3,000 samples for a total sample size of 9,000. Each sample had 300 informative features per modality that distinguished the classes as well as an additional 1800 uninformative features per modality which served as random noise. Of the informative features, 30% were simulated to be modality-shared, with a correlation of 0.3 between corresponding features of different modalities; the remaining features had a correlation of 0 between modalities, representing modality-specific information. To simulate missingness, masks were generated under various missingness scenarios to indicate which samples had each modality available. More details on synthetic data generation are available in Section 5.6.1.

After generating representations using each of the embedding methods, an XGBoost Classifier model was trained to predict the class labels. Performance was evaluated via balanced accuracy, AUROC, and AP. These simulation experiments enabled us to determine under which settings JASMINE outperforms the other methods.

#### 2.3.1 JASMINE is particularly suited for handling a high degree of missing modalities and unbalanced missingness patterns

We first tested the effect of varying the level of overall modality missingness in the four-modality setting. The proportion of missing modalities in the dataset was varied among five settings: 0, 0.38, 0.53, 0.66, and 0.74. This corresponded to varying the proportions of samples with 1, 2, 3, and 4 modalities available, starting with all samples having all modalities available, and gradually increasing the proportion of samples with fewer modalities. The specific proportions of samples with each number of modalities available are detailed in Section 5.6.1. In all of these settings, missingness was balanced in that all modalities were equally likely to be missing for all samples. The balanced missingness results are shown in Figure 2A. At higher missingness rates of 0.64, 0.66 and 0.74, JASMINE outperformed all other methods on all three performance metrics. At low missingness rates, JASMINE slightly underperformed MVAE and CLUE, but as missingness rate increased, MVAE and CLUE showed a more considerable decline in performance. JASMINE, MVAE, and CLUE had the top performance in all cases. DCCA and GCCA, which do not handle missing data, outperformed IntegrAO when no samples had missing modalities, but at a missingness rate of 0.38 and higher, IntegrAO had better performance, likely due to its ability to incorporate incomplete samples. Since the original data, DCCA, and GCCA could not handle incomplete modalities, results were not reported for the highest missingness rate due to insufficient complete samples.

**Fig. 2.**
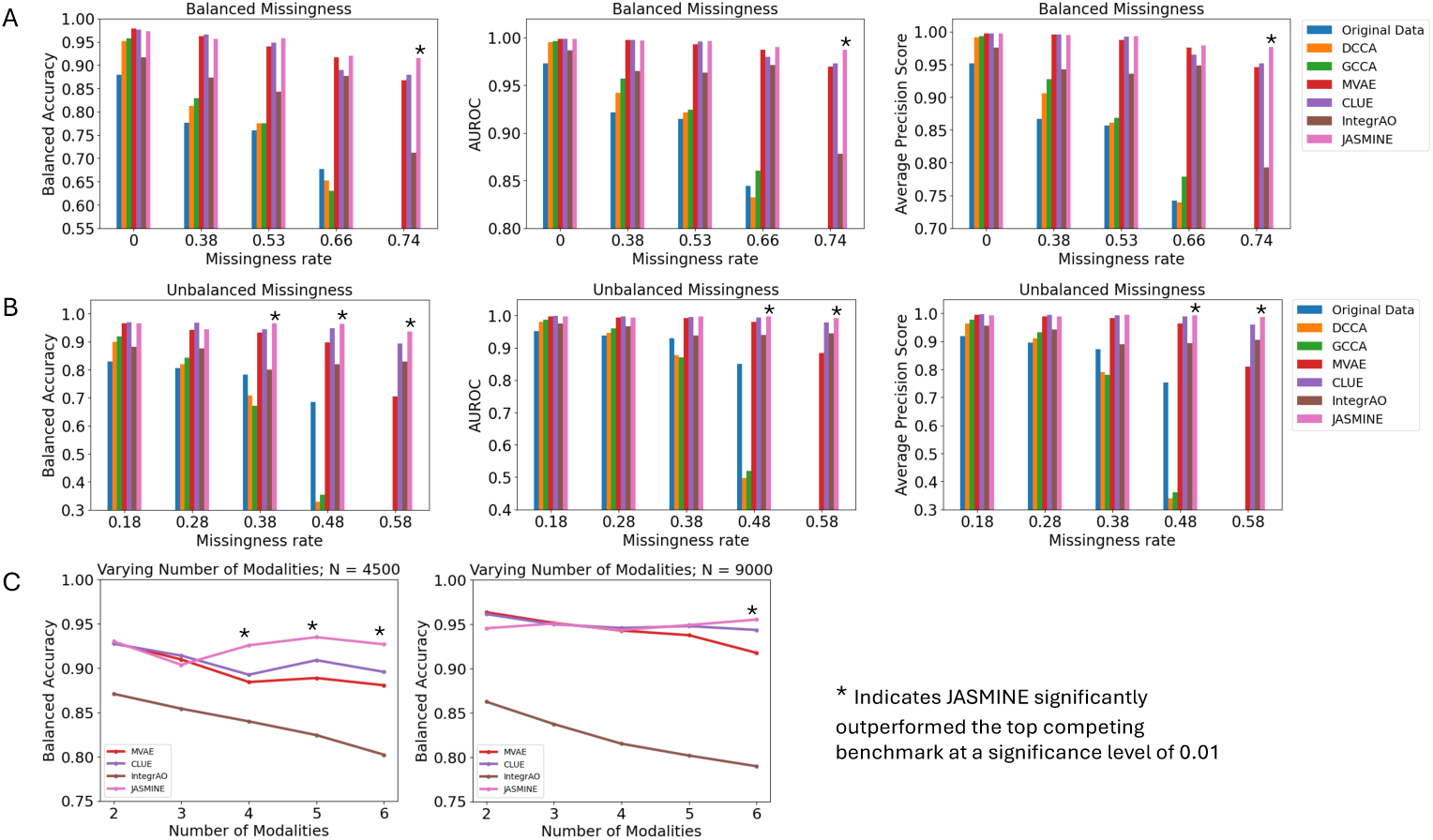
Simulation experiment results. (A) Balanced missingness setting. Overall missingness rate was varied from 0 to 0.74. Probability that a modality was missing was the same for all samples and all modalities. For rate 0.74, methods that do not handle missing data could not be applied due to insufficient samples with all modalities. (B) Unbalanced missingness setting. Missingness patterns differed based on true label, where a different modality was more likely to be missing in each class. Overall missingness rate was varied from 0.18 to 0.58. (C) Varying number of modalities. Sparse modalities were added one-by-one, with each additional modality having a 95% probability of being missing. The values plotted in all graphs represent mean performance over five replicates. Asterisk indicates that JASMINE had significantly better (higher) performance than the second best baseline method at a significance level of 0.01.

Of note, for higher missingness rates, the proportion of samples with one modality is high whereas there is only a small proportion of samples with all modalities available and a moderate proportion of incomplete samples with more than one available. Even when there is a high fraction of samples with one modality, this differs importantly from the unimodal case since the samples in dataset collectively consist of multiple modalities. Thus, methods still need to account for the fact that there are multiple possible modalities and that the specific subset of modalities that are available varies from sample to sample. JASMINE infers missing modality information via the cross-encoders to deduce multimodal information, regardless of how many modalities are available for a given sample. According to our results, JASMINE does this significantly better than the other methods when there is a high degree of missingness.

We also evaluated the performance of these methods when missingness was unbalanced across the classes in the four-modality setting to determine whether systematic class-based differences in modality missingness patterns would impact the embeddings and their performance on downstream classification. In the unbalanced missingness case, the probability of missing a particular modality varied based on a sample’s true label. For example, samples in class 1 were more likely to be missing the first modality, samples in class 2 were more likely to be missing the second modality, and samples in class 3 were more likely to be missing the third modality. Let us denote *p* as the missingness rate when a modality is ‘more likely’ to be missing and *q* as the missingness rate otherwise. We varied *p* and *q* to produce a range of overall modality missingness rates: 0.18, 0.28, 0.38, 0.48, and 0.58. The specific values of *p* and *q* for each setting are detailed in Section 5.6.1. The unbalanced missingness results shown in Figure 2B demonstrate that JASMINE achieves the best classification performance for overall missingness rates of 0.38, 0.48, and 0.58 while maintaining performance on par with the top methods for the lower missingness rates. As with the balanced missingness results, this suggests that JASMINE is particularly advantageous when the data have higher levels of sparsity. Notably, we restricted the test samples to those which were complete. The fact that JASMINE achieves superior performance when tested only on the complete samples suggests that its top performance is not solely due to learning class-specific modality missingness patterns. After JASMINE, CLUE and MVAE performed well, followed by IntegrAO. DCCA and GCCA had a notable decline in performance as missingness rate increased, demonstrating much worse performance than the original data for missingness rates of 0.38 and 0.48. This decline was less pronounced in the balanced missingness setting in Figure 2A, suggesting that these methods struggle more to handle the higher level of class imbalance in the complete samples resulting from unbalanced missingness patterns.

In addition to varying the modality missingness, we investigated the effect of adding sparse modalities for large and small sample sizes. Here, we started with all samples having the first modality, and then iteratively added additional modalities with a 95% probability of being missing. The results are shown in Figure 2C. At both sample sizes, performance showed a notable decline for MVAE, CLUE, and IntegrAO as more modalities were included, whereas it remained more stable for JASMINE. At the smaller sample size of 4,500, this difference in performance was more notable, and JASMINE achieved better performance for 4, 5, and 6 modalities. These patterns were also observed at the larger sample size of 9,000, but performance was more similar among MVAE, CLUE and JASMINE. JASMINE still had significantly better performance when there were 6 modalities. These results demonstrate that with an increasing number of sparsely available modalities, JASMINE achieves significantly better performance than the other methods, demonstrating its ability to better account for multimodal information when arbitrary subsets of the modalities are available. The ability to account for samples with only a subset of modalities becomes increasingly important as more modalities are incorporated, as it becomes more likely that not all samples are complete.

Overall, these results suggest that JASMINE has particularly strong performance relative to the other methods when modalities have a higher degree of missingness and when the number of sparse modalities is higher, especially when sample size is limited. Additionally, JASMINE was shown to be able to handle unbalanced modality missingness better than the other methods at higher overall missingness rates.

### 2.4 Evaluation of representations learned from TCGA multi-omics data

We applied our methods to mRNA-seq, methylation, miRNA-seq, and functional proteomics reverse phase protein array (RPPA) data from The Cancer Genome Atlas (TCGA) [29]. This dataset comprises 30 cancer types across 10,472 total samples, several of which are missing one or more modalities. Missingness is blockwise, such that either all features or no features are missing for a given modality. JASMINE serves not only as a representation learning technique, but also a means of dimensionality reduction. Therefore, to test our methods’ ability to handle a large number of features, we did not perform any prior feature selection or dimensionality reduction. The summary of sample sizes, dimensionality, and labels are available in Table 2A, and more details about TCGA data processing are in Section 5.6.2.

**Table 2.**
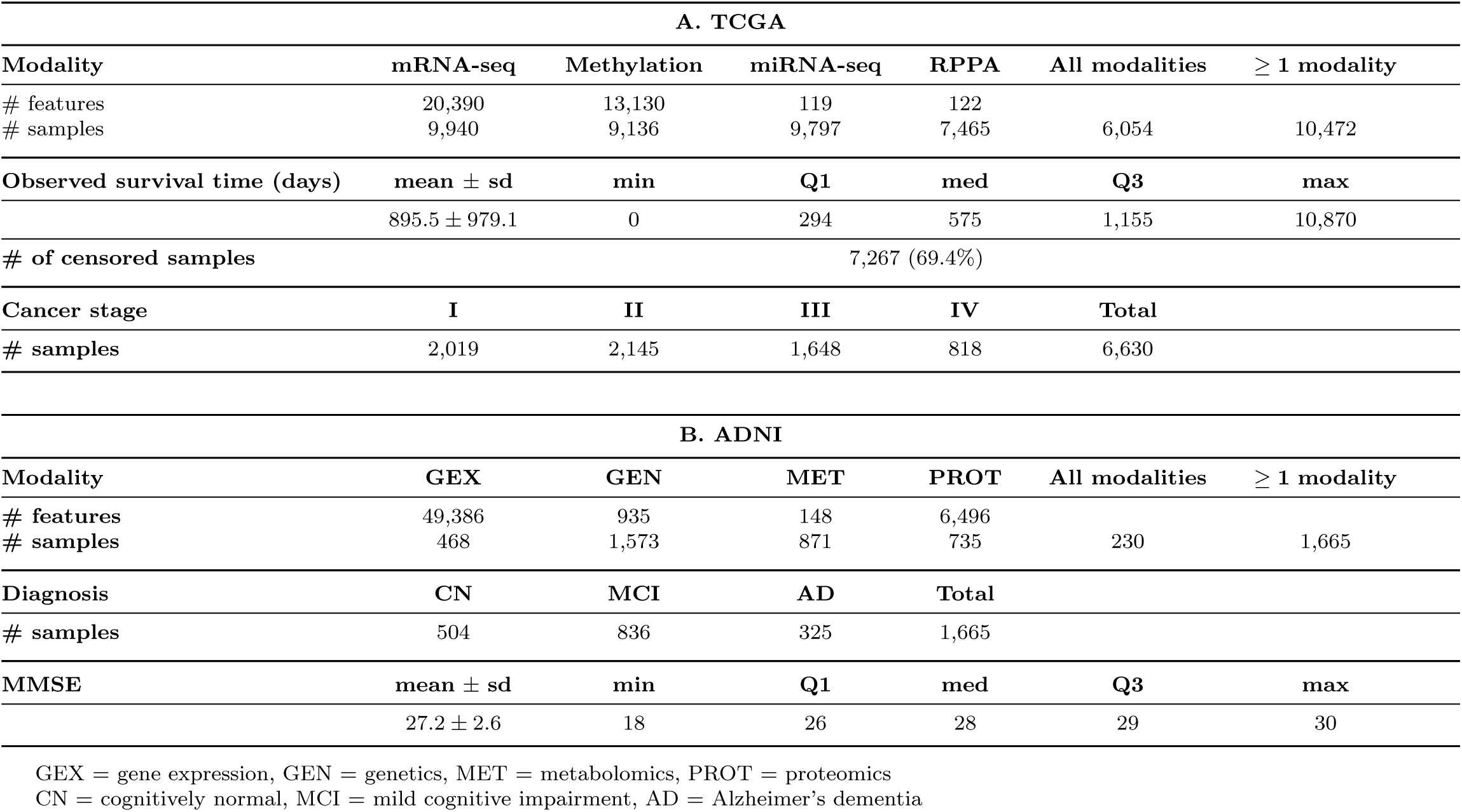
Data Summary.

| A. TCGA |  |  |  |  |  |  |
| --- | --- | --- | --- | --- | --- | --- |
| Modality | mRNA-seq | Methylation | miRNA-seq | RPPA | All modalities | $\geq 1$ modality |
| # features | 20,390 | 13,130 | 119 | 122 |  |  |
| # samples | 9,940 | 9,136 | 9,797 | 7,465 | 6,054 | 10,472 |
| Observed survival time (days) | mean ± sd | min | Q1 | med | Q3 | max |
| | 895.5 ± 979.1 | 0 | 294 | 575 | 1,155 | 10,870 |
| # of censored samples | 7,267 (69.4%) |  |  |  |  |  |
| Cancer stage | I | II | III | IV | Total |  |
| # samples | 2,019 | 2,145 | 1,648 | 818 | 6,630 |  |
| B. ADNI |  |  |  |  |  |  |
| Modality | GEX | GEN | MET | PROT | All modalities | $\geq 1$ modality |
| # features | 49,386 | 935 | 148 | 6,496 |  |  |
| # samples | 468 | 1,573 | 871 | 735 | 230 | 1,665 |
| Diagnosis | CN | MCI | AD | Total |  |  |
| # samples | 504 | 836 | 325 | 1,665 |  |  |
| MMSE | mean ± sd | min | Q1 | med | Q3 | max |
| | 27.2 ± 2.6 | 18 | 26 | 28 | 29 | 30 |

GEX = gene expression, GEN = genetics, MET = metabolomics, PROT = proteomics CN = cognitively normal, MCI = mild cognitive impairment, AD = Alzheimer's dementia

We apply the embeddings generated by our method to two supervised tasks: survival prediction and cancer stage classification. Both are of practical significance: being able to better predict a cancer patient’s survival can help with treatment and palliative care decisions as well as future planning for the patient and their loved ones. Cancer stage classification can help determine the severity of a patient’s condition, aiding in prognosis and treatment decision-making. The learned representations can also be used for unsupervised tasks such as clustering to identify cancer subtypes. To assess their discriminative capabilities, we evaluate their ability to form clusters of patients with differential survival.

#### 2.4.1 JASMINE-generated representations achieved superior performance on survival prediction and cancer stage classification in cancer patients

We used the representations learned by our method to train a Coxnet regression model for survival prediction. The results are shown in Table 3A. JASMINE achieved superior c-index IPCW (0.762) and CD AUC (0.798) compared to the baselines, with a 6.6% improvement in c-index and 3% improvement in CD AUC when compared to the concatenated original data features. Notably, the CCA-based methods had the worst performance, followed by IntegrAO and MVAE, and all four were out-performed by the original data. CLUE performed well, indicating the advantage of preserving modality-specific information. JASMINE displayed further performance gains over CLUE, suggesting that preserving joint and specific information while using contrastive learning and orthogonality constraints is beneficial.

**Table 3.** Results on TCGA multi-omics data.

| **A. Survival prediction results** |  |  |  |
| --- | --- | --- | --- |
| Method | C-index IPCW | CD AUC |  |
| Original Data | $0.717 \pm 0.028$ | $0.774 \pm 0.014$ | |
| DCCA | $0.565 \pm 0.139$ | $0.623 \pm 0.133$ | |
| GCCA | $0.529 \pm 0.131$ | $0.525 \pm 0.146$ | |
| MVAE | $0.692 \pm 0.013$ | $0.749 \pm 0.008$ | |
| CLUE | $0.739 \pm 0.019$ | $0.796 \pm 0.005$ | |
| IntegrAO | $0.652 \pm 0.016$ | $0.673 \pm 0.019$ | |
| JASMINE | **$0.762 \pm 0.013$** | **$0.798 \pm 0.005$** |  |
| **B. Cancer stage classification results** |  |  |  |
| Method | Balanced Acc | AUROC | AP |
| Original Data | $0.415 \pm 0.018$ | $0.683 \pm 0.011$ | $0.438 \pm 0.008$ |
| DCCA | $0.459 \pm 0.013$ | $0.723 \pm 0.006$ | $0.487 \pm 0.008$ |
| GCCA | $0.493 \pm 0.004$ | $0.733 \pm 0.005$ | $0.506 \pm 0.009$ |
| MVAE | $0.469 \pm 0.012$ | $0.728 \pm 0.006$ | $0.498 \pm 0.011$ |
| CLUE | $0.488 \pm 0.011$ | $0.742 \pm 0.006$ | $0.518 \pm 0.009$ |
| IntegrAO | $0.444 \pm 0.010$ | $0.703 \pm 0.008$ | $0.461 \pm 0.015$ |
| JASMINE | **$0.497 \pm 0.007$** | **$0.743 \pm 0.004$** | **$0.521 \pm 0.008$** |

C-index IPCW is the C-index that accounts for high proportion of censoring in the data. CD AUC is the cumulative dynamic AUC for predicting mortality across a range of thresholds. Results are presented as mean ± sd.
AP is average precision, which summarizes a precision-recall curve as the weighted sum of precisions achieved at multiple thresholds. As this is multi-class classification, AUROC and AP are macro-averaged over each class in a one-vs-rest approach. Results are presented as mean ± sd.
Bold font indicates the best performance.

We also used the learned representations as input to a XGBoost classifier to predict cancer stage as I, II, III or IV (see Section 5.6.2 for more details). The results are shown in Table 3B. JASMINE again showed superior balanced accuracy (0.497), AUROC (0.743), and average precision score (0.521). These results suggest that the task-agnostic integrated representations learned by our method achieve the goal of obtaining superior performance across diverse tasks. A further advantage of our method is that it only needs to be trained once on the data and then generates one set of representations that can be utilized for multiple different downstream tasks.

#### 2.4.2 Clustering JASMINE-derived representations suggests biologically meaningful cancer subtypes

In addition to supervised tasks, the representations learned by JASMINE can be used for unsupervised tasks such as clustering. This highlights another advantage to an unsupervised representation learning approach, which does not bias the representations to retain discriminatory information specific to a single task. We investigated the potential for JASMINE-derived embeddings to stratify patients with a given cancer type into biological meaningful subtypes. We applied K-means clustering to the embeddings for five cancer types: breast invasive carcinoma (BRCA), kidney renal clear cell carcinoma (KIRC), lung adenocarcinoma (LUAD), skin cutaneous melanoma (SKCM), and colon adenocarcinoma (COAD) to produce 3-8 clusters. We then compared the p-value of the log-rank test for differential survival among the clusters between JASMINE and the baseline embedding methods. The results are shown in Figure 3. Across cancer types and cluster numbers, clusters derived from JASMINE embeddings more consistently had a high level of survival differentiation compared to the other methods. It often was in the top three methods with highest survival differentiation. This indicates that the JASMINE-derived embeddings preserve information that may be useful for distinguishing cancer subtypes.

**Fig. 3.**
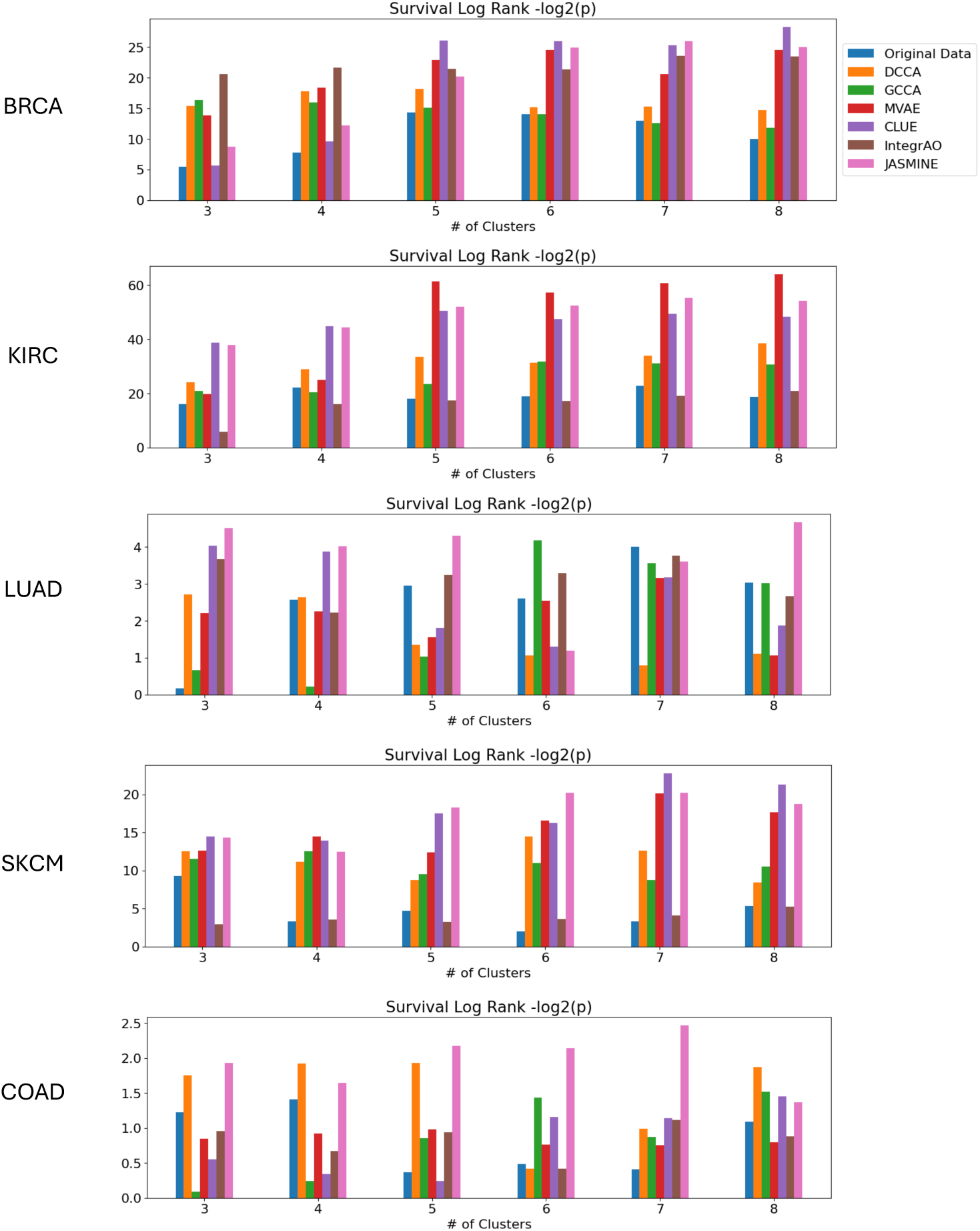
Clustering Analysis. Log-rank test for differential survival among 3-8 clusters in five cancer types in TCGA: breast invasive carcinoma (BRCA), kidney renal clear cell carcinoma (KIRC), lung adenocarcinoma (LUAD), skin cutaneous melanoma (SKCM), and colon adenocarcinoma (COAD). Reported values are mean *−* log_2_(*p*) over five replicates, where *p* is the p-value of the log-rank test.

#### 2.4.3 The most important embedding feature for cancer survival prediction can be explained by known disease markers

We hypothesize that the JASMINE-derived embeddings attain superior performance by capturing important multi-modal information that effectively distinguishes different subjects. Although the embedding features are not directly interpretable, we performed additional analyses to identify which original features from each modality contributed most to the embedding. JASMINE itself is a self-supervised representation learning method, and thus the embeddings it generates are not specialized for a specific task. Thus, employment of JASMINE for a particular prediction problem utilizes a two-step pipeline: first generating the embeddings and then inputting the representations to a downstream prediction model. To identify the features in each modality driving a particular prediction, we also approach feature interpretation in a two-step manner.

The first step involves using the downstream prediction model to identify which embedding features are most important for the prediction task. For this experiment, we focused on the cancer survival prediction task. The downstream predictive model for survival prediction was the Coxnet regression model. We selected the embedding feature corresponding to the regression coefficient with the largest absolute value as the most predictive feature. In our experiment, this was the 28th feature of the mRNAseq-specific embedding (mRNAseq28).

In the second step, we interpret the embedding feature by identifying which original interpretable variables (e.g. genes, miRNAs) from each modality contribute most to those embedding features. We do this in two main steps. First, we cluster the features within each modality into groups of correlated features. Then, for each group, we shuffle the values of all its constituent features and measure the impact on the embedding feature. We identify and select the feature group in each modality that produces the greatest change in the embedding. We expect that these selected interpretable input features are important drivers of the embedding feature of interest and thus can be used to provide a biological explanation of the information it contains. More details on our procedure for identifying the most important input features can be found in Section 5.5.

Applying this approach to the samples from the testing set, we identified top relevant feature clusters for each modality. For mRNAseq, Methylation, and RPPA, the features corresponded to genes, whereas the miRNAseq features corresponded to miRNA molecules. For the top mRNAseq, Methylation, and RPPA gene clusters, we used PANTHER [30] for GO Enrichment Analysis from the Gene Ontology Consortium [31, 32] to determine the top Reactome [33] pathways represented by these genes. For the top miRNA cluster, we used the miRNA Enrichment Analysis and Annotation Tool (miEAA) [34] for over-representation analysis to identify the top associated diseases from the Human microRNA Disease Database (HMDD) [35]. The top enrichments are shown in Table 4.

**Table 4.** Top enriched pathways and diseases associated with the most predictive embedding feature for TCGA survival regression.

| Modality | Reactome Pathway /<br>HMDD disease | FDR-adjusted p-value |
| --- | --- | --- |
| mRNAseq | GABA synthesis | 1.06E-02 |
|  | ↳ GABA synthesis, release, reuptake and degradation | 2.42E-10 |
|  | ↳ Neurotransmitter release cycle | 3.16E-15 |
|  | ↳ Transmission across Chemical Synapses | 7.17E-29 |
| Methylation | Negative feedback regulation of MAPK pathway | 2.92E-04 |
|  | ↳ Negative regulation of MAPK pathway | 1.25E-03 |
|  | Signaling by TCF7L2 mutants | 2.36E-02 |
|  | ↳ Signaling by WNT in cancer | 2.34E-03 |
|  | ↳ Diseases of signal transduction by growth factor<br>receptors and second messengers | 1.39E-11 |
|  | Signaling by MAP2K mutants | 5.84E-03 |
|  | ↳ Oncogenic MAPK signaling | 6.60E-04 |
| miRNAseq | Colorectal Carcinoma | 2.28E-04 |
|  | Carcinoma, Breast | 9.98E-04 |
|  | Pancreatic Neoplasms | 9.98E-04 |
|  | Amyotrophic Lateral Sclerosis | 9.98E-04 |
|  | Vulvar Squamous Tumor | 1.28E-03 |
|  | Glioblastoma | 1.60E-03 |
|  | Carcinoma, Hepatocellular | 1.71E-03 |
|  | Breast Neoplasms | 2.58E-03 |
| RPPA | positive regulation of interleukin-3 production | 4.23E-02 |
|  | ↳ regulation of gene expression | 1.61E-05 |
|  | ↳ regulation of multicellular organismal process | 6.30E-07 |
|  | ↳ positive regulation of gene expression | 1.77E-04 |
|  | ↳ positive regulation of multicellular organismal process | 4.23E-06 |
|  | lysyl-tRNA aminoacylation | 4.22E-02 |
|  | ↳ macromolecule metabolic process | 8.94E-03 |
|  | ↳ protein metabolic process | 1.03E-02 |

Reactome [33] pathways are organized hierarchically. Indented entries preceded with arrows indicate sub-pathways.

For mRNAseq, methylation, and RPPA, the top identified gene clusters are enriched in pathways involved in cancer. This is most obvious for the genes identified for the methylation modality, which were enriched for negative feedback regulation of the mitogen-activated protein kinase (MAPK) pathway, signaling by TCF7L2 mutants including signaling by Wnt in cancer, and signaling by MAP2K mutants including oncogenic MAPK signaling. It is known that the MAPK cascade is involved in human cancer cell survival, malignancy, and resistance to drug therapy [36]. This pathway is commonly deregulated in cancer cells in many different tumor types [37]. The fact that the top genes from the methylation modality were enriched for the oncogenic MAPK signaling pathway clearly suggests an association with MAPK-related cancer mechanisms supported by prior literature [37–40]. Additionally, TCF7L2 is a Wnt transcription factor that is known to suppress cancer cell invasion, and therefore reduced activity of this gene can increase malignancy [41]. It has also been suggested that TCF7L2 mediates resistance to radiotherapy and chemotherapy via its involvement in the Wnt signaling pathway which is known to be involved in cancer [42].

The genes from mRNA that most contributed to the embedding feature were enriched in the GABA synthesis pathway. Synthesis of GABA, a prominent neurotransmitter, has been found to be upregulated in cancer cells, including in non-nervous tissues, and increased GABA levels are associated with poor prognosis in multiple cancer types [43–49]. Prior literature has suggested that this is due to GABA’s involvement in a cancer-intrinsic pathway that results in increased tumor cell proliferation and immune cell suppression [43]. Furthermore, GABA receptors can modulate cancer proliferation and migration through known mechanisms [43, 49, 50]. Additionally, deficient GABA transaminase, which is an enzyme that breaks down GABA, is associated with poor prognosis in hepatocellular carcinoma [51, 52].

The genes from RPPA that most contributed to the embedding feature were enriched in pathways involving the positive regulation of interleukin-3 (IL-3) production and lysyl-tRNA aminoacylation. Interleukins and their associated cytokines are important components of signaling pathways for innate and adaptive immune cells and thus play a critical role in cancer development and progression [53]. Additionally, it has been found that IL-3 acts as a growth factor in many hematologic neoplasms, and its receptor protein CD123 is overexpressed in multiple hematologic cancers [54, 55]. Furthermore, a high expression of CD123 in acute myeloid leukemia is associated with worse prognosis [56]. The lysyl-tRNA aminoacylation pathway was also overrepresented by the top genes from the RPPA modality. Lysyl-tRNA synthetase is a type of aminoacyl-tRNAsynthetase (ARS) which is responsible for aminoacylation, the process by which a specific amino acid is attached to its corresponding tRNA, and plays an essential role in protein synthesis [57]. ARSs maintain cellular and systemic homeostasis, and overexpression or mutations in ARS-encoding genes have been linked to multiple cancer types, including breast, cervical, colorectal, endometrial, head and neck, lung, liver, ovarian, pancreatic, renal, stomach, thyroid, and urothelial cancer, glioma, and melanoma [57–62]. Among the different types of ARSs, lysyl-tRNA synthetase has a relatively high cancer-associated deregulation, can promote metastasis, and is often highly expressed in various cancer cells [63–65]. It has been found to be significantly upregulated in hepatocellular carcinoma [66], breast cancer [67], colorectal cancer [68], and gastric cancer with high expression of KRS in tumor cells associated with poor prognosis [69].

Finally, in the top selected miRNA feature cluster associated with the most predictive embedding feature, many types of cancer, including colorectal carcinoma, breast carcinoma, pancreatic neoplasms, vulvar squamous tumor, glioblastoma, hepatocellular carcinoma, and breast neoplasms, were over-represented. Thus, these results demonstrate that the top genes and miRNAs contributing to the embedding feature most predictive of cancer survival were enriched in pathways related to cancer proliferation, metastasis, and prognosis for multiple types of cancer. This affirms that JASMINE was able to learn an embedding containing information on important cancer biomarkers, aiding in survival prediction.

#### 2.4.4 The top embedding feature for cancer survival prediction captures biologically meaningful inter-modality correspondence

In addition to assessing for known predictive biomarkers of cancer survival, we evaluated the inter-modality correspondence of the input variables represented by the top embedding feature for survival prediction (mRNAseq28). To do this, we found the cluster of features within each modality that contributed most to mRNAseq28. We then assessed the overlap in information among the selected clusters across modalities.

For mRNAseq, Methylation, and RPPA, we identified the genes within the top feature clusters that were shared by at least two modalities. For miRNAseq, we identified the target genes for each miRNA molecule in the selected feature cluster that overlapped with at least two of the other modalities. The target genes and their target prediction scores for each miRNA were obtained from miRDB [70, 71]. Target prediction score ranges between 50-100, with higher score indicating more confidence in the prediction. The results are shown in Figure 4.

**Fig. 4.**
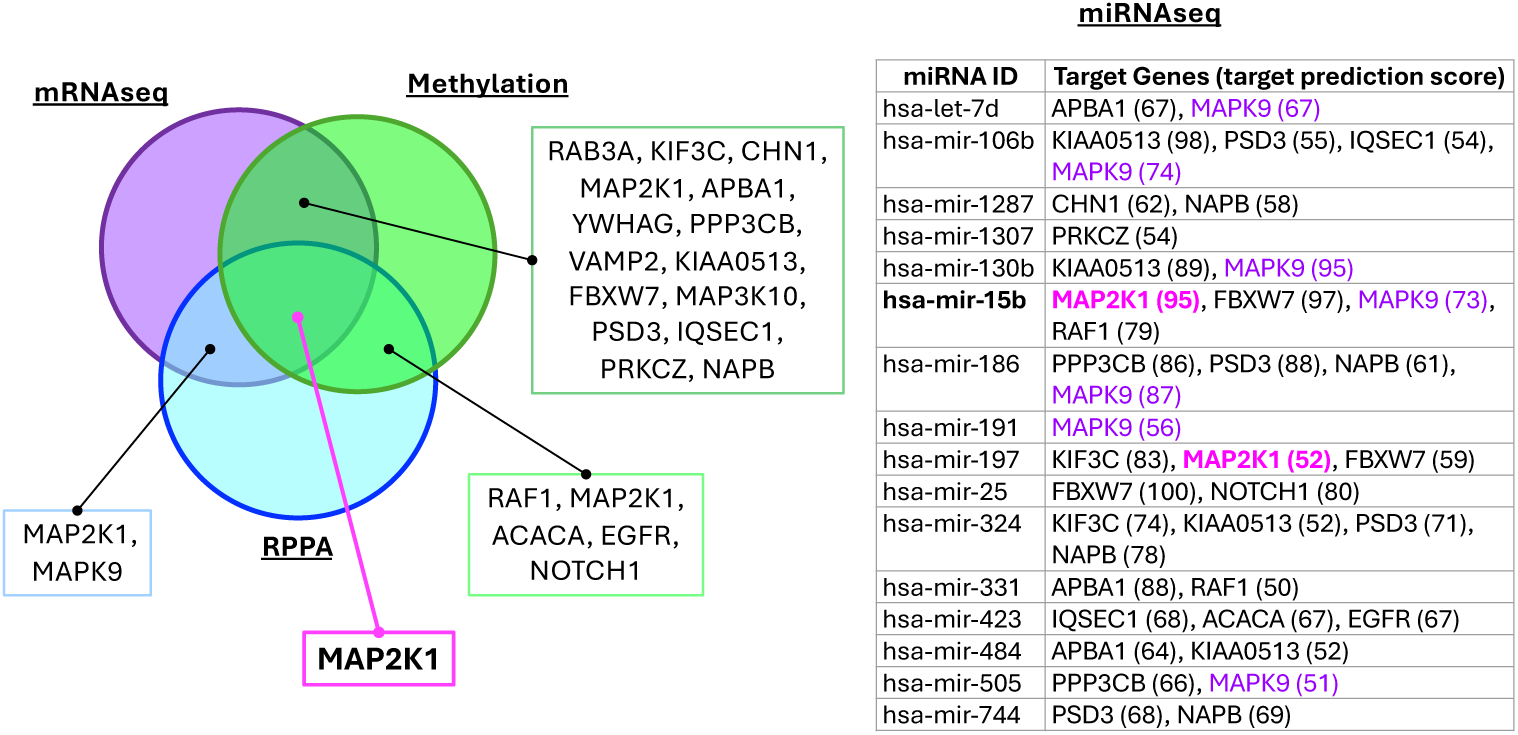
Inter-modal feature correspondence. On the left, the Venn diagram shows the overlapping genes among the top feature clusters for the mRNAseq, Methylation, and RPPA modalities. The miRNA features are miRNA molecule IDs rather than genes; therefore on the right, the miRNA IDs are listed alongside any of their predicted target genes and corresponding target prediction scores in parentheses for any genes that overlapped with at least two of the other modalities. MAP2K1 is colored in pink and bolded as the gene shared by all modalities. MAPK9 is colored in purple as the most frequently targeted gene across the miRNAs in the top miRNAseq cluster. hsa-mir-15b is bolded as the miRNA molecule that targets both MAP2K1 and MAPK9, with high target scores for both.

Among the gene clusters identified for mRNAseq, Methylation, and RPPA, MAP2K1 was found within all three. Two of the miRNAs (hsa-mir-15b and hsamir-197) had MAP2K1 as a predicted target gene. In particular, hsa-mir-15b had a high target prediction score of 95. Gain-of-function mutations in MAP2K1 have been observed in multiple cancers including ovarian [72], melanoma [73], colon [73, 74], and lung [75, 76].

Across all miRNA molecules in the cluster, we also see that MAPK9 was frequently a predicted target. MAPK9 is related to MAP2K1 in its involvement in the MAPK signaling pathway, which regulates cellular processes including proliferation, differentiation, transcriptional regulation, and apoptosis [77, 78]. It has been correlated with poor prognosis and tumor progression in various cancers including glioma [79], prostate cancer [80], breast cancer [78], colorectal cancer [81], and renal cell carcinoma [82]. Again, we note that the target prediction score of hsa-mir-15b for MAPK9 was relatively high, at 73. This may suggest an important role particularly for hsa-mir-15b. The other target genes for hsa-mir-15b that overlapped with at least two of the top mRNAseq, Methylation, and RPPA clusters were FBXW7 (target score = 97) and RAF1 (target score = 79). FBXW7 is a known tumor suppressor gene [83]. It plays a role in the degradation of oncoproteins that are critically involved in oncogenesis, and loss-of-function mutations in this gene are correlated with resistance of tumor cells to treatment and poor prognosis [84–86]. Decreased expression of FBXW7 is observed in multiple cancers, including glioma, lung cancer, liver cancer, urothelial cancer, ovarian cancer, hematopoietic cancers, melanoma, and chronic lymphocytic leukemia [84, 86–90]. RAF1 is a proto-oncogene, and alterations of RAF proteins occurs commonly in cancer [91]. Like MAP2K1 and MAPK9, it is involved in the MAPK signaling pathway, and it has been found to play a role in a variety of solid cancers [92–95].

Thus, the overlap in the most important biological variables across different modalities for the mRNAseq28 embedding feature demonstrated inter-modality correspondence. These consistencies suggest that the cross-encoders that gave rise to the mRNAseq-specific embedding using the information from other modalities were able to capture biologically meaningful relationships among the different modalities. Furthermore, the corresponding variables involved in these relationships were relevant markers for cancer progression and prognosis.

### 2.5 Evaluation of representations learned from ADNI multi-omics data

To demonstrate the applicability of our method to more than one dataset, we also applied JASMINE to gene expression, genetic, metabolomics, and proteomics data from the Alzheimer’s Disease Neuroimaging Initiative (ADNI) [96, 97]. There is a total of 1,665 samples, including several that have at least one missing modality. These data are summarized in Table 2B, and more details about ADNI data processing are in Section 5.6.3.

We applied the representations generated by JASMINE to two tasks: diagnosis classification and cognitive score prediction at baseline. Diagnosis classification is an important problem in Alzheimer’s disease research and treatment. Alzheimer’s disease (AD) is a complex and heterogeneous disease with a wide array of underlying mechanisms and presentations that complicate its characterization and diagnosis. Being able to classify individuals based on their multi-omics data may provide insight into the utility of molecular information for AD detection, guide further research into the molecular signatures underlying AD, and improve diagnostic models. The MMSE (Mini-Mental State Examination) is a commonly used cognitive score which is often used in diagnosing AD. This provides a measure of an individual’s cognitive functioning on a more fine-grained scale than a diagnostic label. Predicting MMSE using multi-omics data may also provide insight into the link between a subject’s molecular profile and cognitive functioning, as well as how these relate to AD progression. By evaluating the representations generated by JASMINE on both classification and regression tasks, we also demonstrate a wider range of prediction problems to which our method can be applied.

#### 2.5.1 JASMINE-generated representations achieved superior performance on AD diagnosis classification and cognitive score prediction

We applied JASMINE-derived representations to binary classification between CN and MCI/AD. The results are shown in Table 5A. JASMINE outperformed all baseline methods on all three classification metrics (balanced accuracy = 0.587, AUROC = 0.659, AP = 0.822), again demonstrating the ability of the proposed method to produce representations that have superior performance on a variety of tasks across different datasets. This also suggests that multi-omics data may have some utility in detecting MCI/AD.

**Table 5.** Results on ADNI multi-omics data.

| **A. Diagnosis classification results** |  |  |  |
| --- | --- | --- | --- |
| Method | Balanced Acc | AUROC | AP |
| Original Data | $0.490 \pm 0.046$ | $0.507 \pm 0.045$ | $0.722 \pm 0.033$ |
| DCCA | $0.484 \pm 0.024$ | $0.504 \pm 0.041$ | $0.706 \pm 0.031$ |
| GCCA | $0.536 \pm 0.061$ | $0.577 \pm 0.044$ | $0.738 \pm 0.039$ |
| MVAE | $0.566 \pm 0.089$ | $0.624 \pm 0.085$ | $0.797 \pm 0.051$ |
| CLUE | $0.571 \pm 0.028$ | $0.613 \pm 0.056$ | $0.765 \pm 0.043$ |
| IntegrAO | $0.498 \pm 0.060$ | $0.439 \pm 0.074$ | $0.662 \pm 0.057$ |
| JASMINE | **$0.587 \pm 0.050$** | **$0.659 \pm 0.044$** | **$0.822 \pm 0.028$** |

| **B. MMSE prediction results** |  |  |  |
| --- | --- | --- | --- |
| Method | RMSE | $R^2$-score | r-pred |
| Original Data | $2.302 \pm 0.134$ | $0.006 \pm 0.118$ | $0.185 \pm 0.214$ |
| DCCA | $2.320 \pm 0.109$ | $-0.009 \pm 0.041$ | **$0.198 \pm 0.105$** |
| GCCA | $2.428 \pm 0.061$ | $-0.103 \pm 0.055$ | $-0.035 \pm 0.110$ |
| MVAE | $2.329 \pm 0.075$ | $-0.015 \pm 0.065$ | $0.143 \pm 0.110$ |
| CLUE | $2.314 \pm 0.050$ | $-0.002 \pm 0.043$ | $0.192 \pm 0.049$ |
| IntegrAO | $2.421 \pm 0.072$ | $-0.096 \pm 0.066$ | $-0.021 \pm 0.150$ |
| JASMINE | **$2.301 \pm 0.049$** | **$0.010 \pm 0.042$** | $0.180 \pm 0.070$ |

AP is average precision, which summarizes a precision-recall curve as the weighted sum of precisions achieved at multiple thresholds. Results are presented as mean ± sd.
$R^2$ score is the coefficient of determination, as computed using the `r2.score` function in `scikit-learn`[98]. r-pred is the Pearson correlation coefficient between the predicted and true values. Results are presented as mean ± sd.
Bold font indicates the best performance.

We also applied the embeddings to MMSE score regression. The results are presented in Table 5B. Our proposed method had the best RMSE (2.301) and *R*^2^-score (0.010), suggesting more accurate predictions. We note that while DCCA and the original data had the first and third highest r-pred, respectively (DCCA r-pred = 0.198; orig data r-pred = 0.185), the variation in this metric is very high. These methods are also disadvantaged by the fact that DCCA only can take two modalities at a time, and neither can handle missing data. While CLUE has a higher r-pred than the proposed method (CLUE r-pred = 0.192; JASMINE r-pred = 0.180), it has a negative *R*^2^ score (CLUE *R*^2^ = −0.002) and worse RMSE (CLUE RMSE = 2.314), indicating that while it can predict relative scores, its predictions are less accurate.

### 2.6 PoE, MCL, SCL, and orthogonality constraints enhanced performance on multiple tasks

We performed ablation experiments to evaluate the utility of the main components of JASMINE: PoE for the joint representation (POE), modality-level contrastive learning (MCL), sample-level contrastive learning (SCL), and orthogonality constraints (orthog). We removed one component at a time and evaluated the performance on each of the supervised tasks for both TCGA and ADNI. The results are shown in Table 6. The full model, consisting of all components, had the best performance across most prediction tasks, including TCGA survival prediction, cancer stage classification, and ADNI diagnosis classification. For survival prediction, although there is a very small increase in CD AUC (+0.63%) when POE is dropped, there is also a large drop in c-index (−1.3%), suggesting that POE is still important for this task. Removing the MCL component of the model led to the largest drop in performance across all TCGA tasks, suggesting that modality-level contrastive learning was particularly important. Removing SCL similarly led to relatively large drops in performance in most cases. MCL helps to separate distinct samples, while SCL helps to enhance the similarity structure of the original data, allowing for better discrimination between individuals with different survival times and cancer stages. For TCGA cancer stage classification, dropping orthogonality constraints led to a small improvement in AUROC (+0.40%) and AP (+0.96%), but a much larger drop in balanced accuracy (−2.0%), suggesting that this model component is still beneficial. Using orthogonality constraints to minimize redundancy between the modality-specific and shared components of the embeddings may better preserve important information, aiding in the separation of samples by cancer stage.

**Table 6.** JASMINE ablation experiment results.

| TCGA Survival Prediction |  |  |  |
| --- | --- | --- | --- |
|  | C-index IPCW | CD AUC |  |
| Full Model | **0.762 ± 0.013** | 0.798 ± 0.005 |  |
| -POE | 0.752 ± 0.016 | **0.804 ± 0.009** |  |
| -MCL | 0.726 ± 0.017 | 0.789 ± 0.007 |  |
| -SCL | 0.738 ± 0.017 | 0.791 ± 0.005 |  |
| -orthog | 0.748 ± 0.024 | 0.798 ± 0.005 |  |
| TCGA Cancer Stage Classification |  |  |  |
|  | Balanced Acc | AUROC | AP |
| Full Model | **0.497 ± 0.007** | 0.743 ± 0.004 | 0.521 ± 0.008 |
| -POE | 0.488 ± 0.008 | 0.745 ± 0.003 | 0.523 ± 0.005 |
| -MCL | 0.477 ± 0.007 | 0.738 ± 0.003 | 0.512 ± 0.008 |
| -SCL | 0.486 ± 0.007 | 0.739 ± 0.006 | 0.516 ± 0.013 |
| -orthog | 0.487 ± 0.009 | **0.746 ± 0.004** | **0.526 ± 0.007** |
| ADNI Diagnosis Classification |  |  |  |
|  | Balanced Acc | AUROC | AP |
| Full Model | **0.587 ± 0.050** | **0.659 ± 0.044** | **0.822 ± 0.028** |
| -POE | 0.566 ± 0.055 | 0.643 ± 0.091 | 0.802 ± 0.065 |
| -MCL | 0.498 ± 0.028 | 0.593 ± 0.050 | 0.790 ± 0.045 |
| -SCL | 0.550 ± 0.054 | 0.611 ± 0.052 | 0.785 ± 0.032 |
| -orthog | 0.518 ± 0.047 | 0.559 ± 0.035 | 0.755 ± 0.025 |
| ADNI MMSE Regression |  |  |  |
| | RMSE | $R^2$-score | r-pred |
| Full Model | 2.301 ± 0.049 | 0.010 ± 0.042 | 0.180 ± 0.070 |
| -POE | 2.391 ± 0.091 | -0.070 ± 0.080 | 0.060 ± 0.135 |
| -MCL | **2.284 ± 0.043** | **0.024 ± 0.037** | **0.225 ± 0.050** |
| -SCL | 2.325 ± 0.094 | -0.012 ± 0.080 | 0.142 ± 0.144 |
| -orthog | 2.303 ± 0.079 | 0.007 ± 0.069 | 0.198 ± 0.088 |

Results are presented as mean ± sd. Bold font indicates the best performance.

On the ADNI dataset, we again see that all components of the model are helpful for achieving higher balanced accuracy, AUROC, and AP for diagnosis classification; removing any of them leads to a considerable drop in performance. For the MMSE prediction task, removing MCL from the model led to the best RMSE (2.284), *R*^2^-score (0.024), and r-pred (0.225), but the full model had second-best RMSE (2.301) and *R*^2^-score (0.024), and removing MCL led to large drops in performance on most other tasks. Overall, the ablation results suggest that all elements of JASMINE are useful across most scenarios, encompassing two different datasets and multiple prediction problems.

### 2.7 JASMINE achieves an advantageous balance of computational resource consumption and performance

We compared the computational resource usage of JASMINE and the baseline embedding methods. The training times and memory usage of each of these methods are shown in Table 7. All methods except for GCCA utilized A100 GPUs, whereas GCCA was run on a CPU. Furthermore, all but DCCA could be run using 3 or fewer cores. We note that while GCCA has the shortest training time, it still has relatively high peak memory usage and often has worse performance on many tasks. MVAE took many more epochs to converge than CLUE and JASMINE, resulting in a much longer training time. IntegrAO had much higher memory usage due to its reliance on building patient similarity networks. Overall, CLUE and JASMINE appear to achieve a balance of moderate training time and peak memory usage while attaining high accuracy across multiple tasks.

**Table 7.**
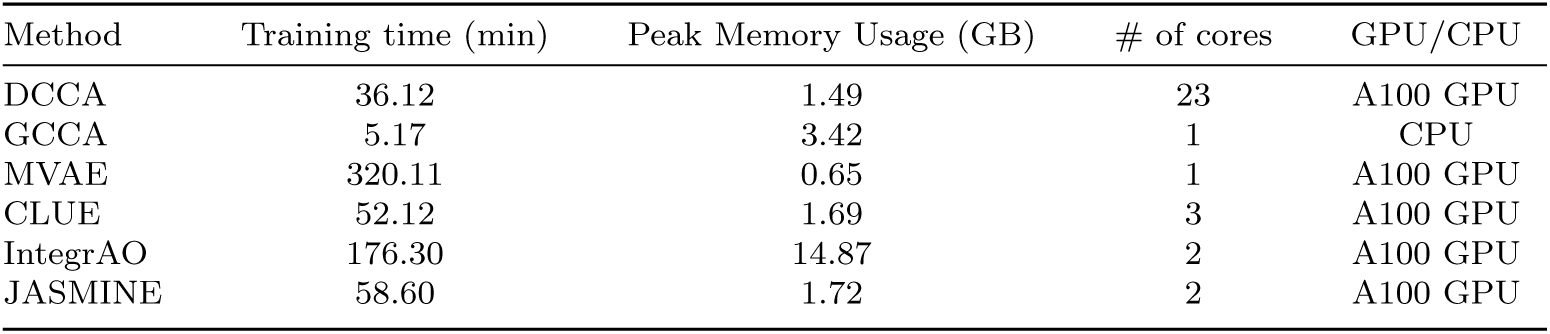
Compute resource comparison.

| Method | Training time (min) | Peak Memory Usage (GB) | # of cores | GPU/CPU |
| --- | --- | --- | --- | --- |
| DCCA | 36.12 | 1.49 | 23 | A100 GPU |
| GCCA | 5.17 | 3.42 | 1 | CPU |
| MVAE | 320.11 | 0.65 | 1 | A100 GPU |
| CLUE | 52.12 | 1.69 | 3 | A100 GPU |
| IntegrAO | 176.30 | 14.87 | 2 | A100 GPU |
| JASMINE | 58.60 | 1.72 | 2 | A100 GPU |

## 3 Discussion

We presented JASMINE, a variational autoencoder-based self-supervised representation learning method that generates task-agnostic embeddings from multi-omics data and is capable of handling samples with arbitrarily missing modalities. The representations are learned to incorporate both modality-specific and shared information while minimizing the redundancy between them. Contrastive learning is used to encourage the representations to enhance the similarities and differences between samples in the embedding space. JASMINE thus addresses the limitations of existing multi-omics integration methods by handling data with arbitrarily missing modalities while preserving both modality-specific and shared information and sample similarity structure. Furthermore, the self-supervised nature of JASMINE enables it to learn representations that preserve information in a data-driven, rather than task-biased, manner. Simulation experiments suggest that JASMINE is particularly well-suited for settings where the rate of modality missingness is high and when the missingness patterns differ among ground truth classes. JASMINE also demonstrates stable performance as additional sparsely available modalities are incorporated, whereas other methods show a notable decline in performance, especially at small sample size. On real data, JASMINE outperforms the baselines across two datasets and multiple prediction tasks, including survival prediction, classification, clustering, and regression. Notably, the three VAE-based methods, MVAE, CLUE, and JASMINE, outperformed the graph-based IntegrAO approach, which tended to surpass the CCA-based methods, on the survival prediction, classification, and regression tasks. Upon further analysis of the JASMINE embeddings of the TCGA data, we found that the embedding feature most important for cancer survival prediction could be explained by known cancer biomarkers that had task-relevant cross-modal correspondence, demonstrating JASMINE’s ability to learn and retain biologically meaningful information. JASMINE’s superior performance across most supervised tasks in addition to its ability to derive biologically meaningful clusters demonstrates the generalizability of its learned embeddings across a wide set of tasks. The ability to use a single set of embeddings that excel at multiple tasks highlights its efficiency as well as its effectiveness at extracting important information from the data.

The self-supervised nature of JASMINE renders it efficient, as it only needs to be trained once per dataset to produce representations with top performance across a variety of tasks. This mechanism also underlies foundation models, which are trained on vast, often unlabeled datasets and can improve performance on downstream tasks [99–101]. We found that JASMINE-derived embeddings had superior performance on a variety of downstream tasks; given large amounts of data, JASMINE could potentially serve as the backbone architecture for an effective foundation model that can fully leverage the potential of incomplete multi-omics data. Additionally, JASMINE could be fine-tuned using specific prediction objectives to generate task-oriented embeddings with potentially even better performance.

Some limitations of our study include limited sample sizes, particularly in the ADNI dataset. Nonetheless, our results show some promise and are expected to improve when more samples are available. Another limitation is that JASMINE models the likelihood function for all modalities as multivariate Gaussians, which may be less ideal for certain data types such as allele counts. Expanding JASMINE to enable the decoders to model a broader range of likelihood functions is a potential direction for future work. We also recognize that our use of a manual tuning procedure for the weights of the loss components in the objective function, while designed to efficiently find a parameter configuration with approximately optimal performance, may be less than ideal. Another extension of this work could be to implement a more precise data-driven approach. Additionally, while we performed analyses to help understand the biological underpinnings of the JASMINE embeddings, we acknowledge that the model itself is not inherently interpretable, the explainability of the embeddings remains understudied, and enhancing interpretability is a direction for future research. Furthermore, our current formulation does not include a quantitative measure of the modality-specific information contained in the final representations. Such quantification utilizing mathematical frameworks like those described in [102], [103], and [104] could be useful for increasing the interpretability of our representations.

JASMINE is also limited to tabular data inputs. In this paper, we focused on applying JASMINE to multi-omics data, which have this format. However, JASMINE could potentially be applied to tabular data from other domains. Furthermore, non-tabular data types, such as images or text, could be vectorized via a preprocessing step and then used as input to our method. In this way, JASMINE could be applied to other types of data from additional disciplines in a relatively straightforward manner. However, utilizing an ad-hoc preprocessing step to vectorize non-tabular modalities may not fully leverage the rich information contained in these data types. Thus, some future directions include extending the model architecture to handle additional modalities such as imaging, spatial transcriptomics, temporal data, and text.

## 4 Conclusions

We introduced JASMINE, a novel self-supervised representation learning method that produces integrated, low-dimensional embeddings of incomplete multi-omics data that better preserve modality-specific and shared information as well as sample similarity structure for enhanced performance across a variety of tasks. JASMINE needs to be trained only once per dataset to produce representations with superior performance in multiple scenarios. Simulation studies demonstrated JASMINE’s advantage in handling sparse data and varying missingness patterns. Upon applying JASMINE to incomplete multi-omics data from two different landmark databases for cancer and Alzheimer’s disease, JASMINE produced representations with superior performance across a range of supervised tasks and which clustered into cancer subtypes with better survival differentiation compared to state-of-the-art baseline multi-omics embedding methods. These results exhibit JASMINE’s ability to better capture useful information for both clinically relevant prediction tasks and subtype clustering. Its demonstrated information extraction capability also suggests that it could potentially serve as the backbone for an effective foundation model that can harness the full potential of incomplete multi-omics data. This has several implications for the future of biomedical research and clinical care, from enhancing the study of underlying disease mechanisms to improved diagnosis and prognosis.

## 5 Methods

### 5.1 Notation and problem

Suppose the input data consists of *n* samples and *m* modalities. Then, we can represent the entire input data as a list of the data matrices for each modality: *X* = *X*^(1)^*, X*^(2)^*, …, X*^(*m*)^ . It is possible that some samples are missing a subset of modalities. Thus, the dimensions of each of these modality-specific data matrices can vary. For example, the data matrix for modality *j*, which is measured for *n_j_* samples and consists of *d_j_* features, has dimension 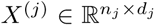 . The goal of our method is to learn integrated representations of the data, preserving both modality-specific and modality-shared information while also allowing for missing modalities. To preserve the most discriminative information in the data while reducing the data dimensionality and avoiding biasing the representations toward a specific downstream task, these representations are learned in a self-supervised manner. These clean and compact representations can then be used as input for a variety of downstream tasks, including regression and classification.

### 5.2 Technical details of JASMINE

To accomplish these goals, we propose a variational autoencoder (VAE)-based model to produce integrated embeddings that comprise both modality-specific and modality-shared, or joint, components. The model architecture consists of the following units:

- *m* self-encoders, 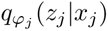, which map a modality *j* to its own latent space to generate latent embedding *z_j_*.
- *m*(*m* − 1) cross-encoders, 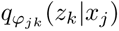, which map a modality *j* to the latent space of another modality *k* to generate latent embedding *z_k_*.
- *m* decoders, 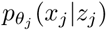 which map a latent embedding for modality *j* back to its original feature space.

Self-encoders are similar to encoders in a vanilla VAE, enabling the latent representation for a given modality to be derived from the modality itself. However, only having self-encoders is insufficient when there are missing modalities. Therefore, the model also consists of cross-encoders which learn to take one modality *j* as input and infer the latent space of another modality *k*. All modality pairings are considered by this model, such that there are *m*(*m* − 1) cross-encoders. This not only allows the method to generate representations for missing modalities using the modalities that are available, but it also enables the learning of inter-modality relationships.

Additionally, there are decoders for every modality, which takes an embedding from the latent space of a given modality as input, and outputs the parameters of the distribution in the original feature space for that modality. The decoders are used for reconstructing the input during training to ensure that the learned embeddings preserve enough information to be able to recover the original data (measured via reconstruction loss). For a visual overview of JASMINE, a schematic is given in Figure 1.

The encoders and decoders are feedforward neural networks with a chosen number of layers and hidden dimensions. In our experiments, the encoders had 2 layers, with hidden dimension of 512. The decoder had 1 layer with hidden dimension of 256. The latent representations have dimension 64.

To learn the parameters of these inference networks, we optimize the following objective function, consisting of the evidence lower bound (ELBO) loss, as is traditionally used for training VAEs:

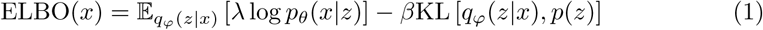

where *p*(*z*) is the standard Normal prior. Here, we see that the first term seeks to maximize the log likelihood of the data given the latent representation, and is equivalent to minimizing the reconstruction loss between the original data and its reconstruction from its embedding. For the second term, KL is the Kullback-Leibler divergence and is used as a regularization mechanism by encouraging the inferred latent distribution to be close to the prior. *λ* and *β* are weights that control the tradeoff between the two terms. For simplicity, JASMINE uses a multivariate Gaussian for the likelihood function in the ELBO for all data modalities.

There is an ELBO term for each of the *m*^2^ encoders, where the first term is calculated using each of the modality-specific decoders to reconstruct the modality for which the latent representation *z* was generated. The second term is calculated as the KL divergence between each of the *m*^2^ inferences and the prior. Thus, we have

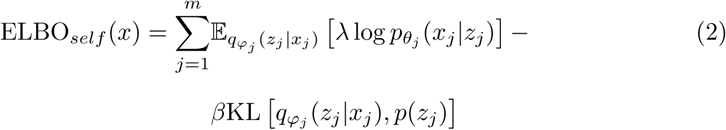

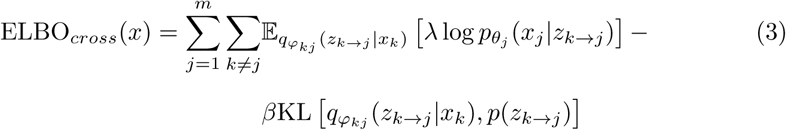

Since an embedding space of a given modality can be generated using any of the *m* modalities via the self- or cross encoders, it is important that these embeddings are consistent regardless of which modality was used to for inference. To do this, we also include a cosine similarity loss term, which ensures that the inferences are aligned in the embedding space. This is given by

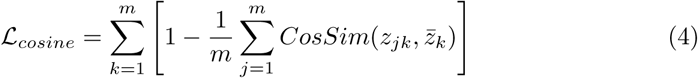

where *z_jk_* is the latent embedding of modality *k* inferred from modality *j*, *z̅_k_* is the mean of *z_jk_* across all *j* ∈ [1*, …, m*] and 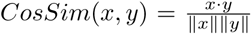. This aims to maximize the similarity between the embeddings inferred by each of the modalities and their mean.

The embeddings given by the self- and cross-encoders contain modality-specific information. In addition to this, the proposed method identifies information that is consistent among all modalities, which is encoded in a joint embedding and composes the modality-shared component of the final learned representation.

#### 5.2.1 Joint embedding via product-of-experts

Information that is shared among all modalities is also important for defining a given sample. Thus, in addition to modality-specific information obtained via the self- and cross-encoders, the proposed method learns a joint embedding to capture information that is shared across the modalities.

The strategy used is product-of-experts (PoE)[13], which defines the joint latent distribution to be the product of the modality-specific latent distributions:

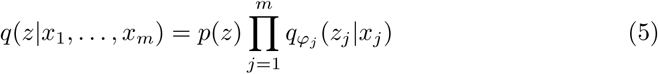

where *p*(*z*) is the standard normal prior and *q*(*z*|*x*_1_*, …, x_m_*) is the joint latent space across all modalities *j* = 1*, …, m*. This strategy handles missing modalities by taking the product over the subset of modalities that are available for a given sample. Let *M_i_* be the set of modalities available for sample *i*. Let us represent sample *i* as *X_i_*, where the data of the *j*th modality for *X_i_* is represented as *x_ij_*. Then, the joint latent representation is estimated as

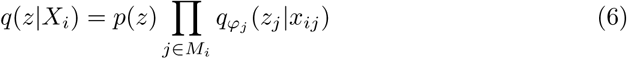

In order to avoid having to learn all 2*^m^*VAEs to account for all possible combinations of modalities, prior methods have approached this estimation by randomly masking subsets of the modalities in the training data. Since our training data already consists of samples with arbitrarily missing modalities, the model can be trained on these samples to learn to infer the joint space in the presence of incomplete modalities. We define an ELBO loss term for the joint inference as follows:

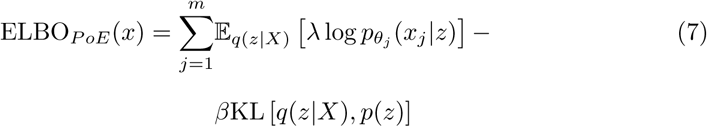

where *z* is the PoE-derived joint representation, *X* refers to all modalities available for a given sample (which may only be a subset of all *m* modalities), *q*(*z*|*X*) is the joint multimodal inference, and *p*(*z*) is the standard normal prior for the joint latent space. Note that the first term is equivalent to minimizing the reconstruction loss of each modality *j* from the joint representation using the corresponding modality-specific decoder, 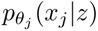. Since PoE produces a single joint embedding that preserves sufficient information to reconstruct each modality, this training process encourages the joint embedding to contain information that is shared among all modalities.

One of the benefits of the PoE-based formulation of the joint representation space is that the resulting joint distribution can be written in closed-form. Namely, given that we represent each modality-specific embedding space as a Normal(*µ_j_*,Σ*_j_*) distribution, the resulting joint latent space is Normal(*µ_PoE_*,Σ*_PoE_*)[105], where:

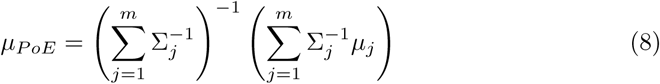

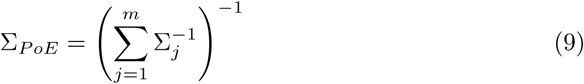

#### 5.2.2 Construction of the final representation

Once the model has obtained the modality-specific representations from the self- and cross-encoders, we obtain final modality-specific representations for each sample by averaging over the representations derived from every available modality, i.e., letting *z_ij_* be the final modality-specific representation of modality *j* for sample *i*, *z_i,k_*_→*j*_ be the representation of modality *j*derived from modality *k*, and *M_i_* be the set of modalities available for sample *i*, we have

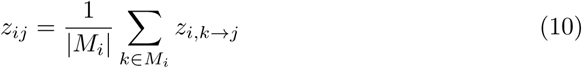

This enables the generation of modality-specific embeddings for all modalities for every sample, even when the modality itself is missing.

Let *z_i,joint_* be the joint representation for modality *i*. Then, the final JASMINE representation for sample *i*, *z_i_*, is the concatenation of each of the modality-specific representations and the joint representation:

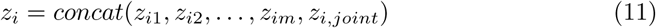

Letting *h* be the dimension of the latent embeddings, it follows that *z_i_* ∈ ℝ*^h^*^(^*^m^*^+1)^.

#### 5.2.3 Orthogonality Constraints

Another aim of our representations is to preserve both modality-specific and modality-shared information in the data while minimizing the amount of redundancy between them, in order to produce more information-rich representations. To do this, we also introduce an orthogonality constraint [18] into our model as a loss term. The orthogonality constraint-based loss term is calculated as follows:

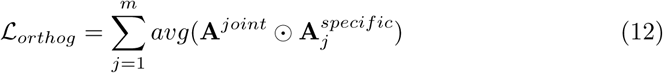

where **A** represent *B* × *h* matrices of the latent representations, *B* represents the batch size, *h* represents the dimension of the joint and specific embeddings, and ⊙ represents element-wise multiplication. The average of this element-wise product provides a measure of the similarity between the joint and modality-specific embeddings, and therefore, minimizing this helps to accomplish our goal of ensuring that the modality-specific embeddings contain different information from the modality-shared embeddings. This helps to maximize the amount of unique information that is preserved in the final representations, which consist of the concatenation of the modality-specific and joint representations.

#### 5.2.4 Contrastive learning

To better enforce consistency among modalities for a given sample as well as preserve and enhance the structure in the original data for better discriminability in down-stream tasks, we introduce two types of contrastive learning (CL), which we refer to as modality-level and sample-level. These losses are applied to the embeddings generated by our model.

Modality-level CL (MCL) promotes similarity between embeddings of modalities from the same sample and difference between embeddings of modalities from different samples to better distinguish different samples while encouraging consensus among the modalities for a given sample. Here, sample affiliation is used to define positive and negative pairs among the modalities: positive pairs consist of embeddings of different modalities from the same sample, and negative pairs are embeddings of modalities from other samples. As there are multiple positive and negative pairs defined for each embedding, we define a multi-way version of contrastive loss based on soft-nearest neighbors loss to calculate MCL. Letting *z_ij_*represent the embedding of modality *j* for the *i*th sample, and *m* be the number of modalities, the MCL is calculated as:

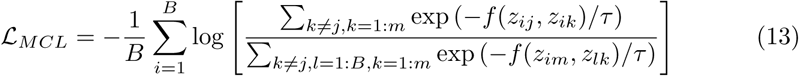

where *B* is the batch size, *f* (*x, y*) is the cosine similarity between *x* and *y*, and *τ* is a temperature parameter that controls how close the features are in the representation space. We set *τ* = 0.1 for all experiments in this study. Note that the numerator consists of the sum of similarity measures over positive pairs, whereas the denominator consists of the sum of similarity measures over all pairs. We seek to maximize the similarity between positive pairs compared to negative pairs. Thus, minimizing the negative log of this ratio accomplishes this goal.

Sample-level CL (SCL) encourages the embeddings to preserve and enhance the similarity structure between samples, based on the joint component of their embeddings. We do this in an unsupervised manner to avoid biasing the representations toward a specific structure based on a single set of labels. Thus, SCL is based on a deep clustering approach [106, 107] that consists of the following main steps: (1) stochastic data augmentation to transform each input data example randomly resulting in two correlated views of the same example, which we denote as a positive pair, and (2) generation of the joint embedding for the augmented dataset (consisting of original and perturbed data), and (3) computation of the contrastive loss on the augmented data, defining the original data and its perturbed representation as a positive pair. We calculate SCL as:

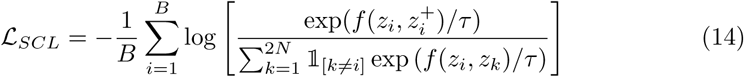

where *z_i_* is the PoE-derived joint component of the embedding, 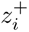 is the augmented version of *z_i_*, and *f* (*x, y*) is again defined to be the cosine similarity between *x* and *y*. Since the data are tabular, the stochastic data augmentation is performed following SCARF[108]: a fraction of the features are randomly selected to be “corrupted” by replacing them with values sampled with replacement from all possible values for that feature within the batch. The goal of SCL is to learn to keep samples that are similar to one another closer together in the embedding space. By defining the original and perturbed versions of each sample as positive pairs, we can accomplish this goal without predefined cluster labels. One of the hyperparameters associated with this loss term is the proportion of features to randomly perturb in each sample. We chose this value to be 0.25, which worked well in our experiments.

#### 5.2.5 Learning objective

Given the loss terms described previously, the final objective function is

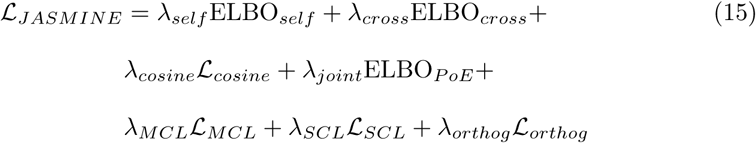

where the *λ* values are scalar weights on the loss terms. These values were tuned manually on the validation data (see Section 5.2.6 for more details on the tuning procedure).

#### 5.2.6 Parameter tuning procedure

The weights of the loss terms of the objective function were tuned manually. We started with a set of values which put each loss term on a similar scale to enable roughly equal contribution to the overall objective. Then, we explored a range of values above and below each initial parameter value, modifying one variable at a time. Based on the direction of change that led to performance improvements on the validation data, we either explored additional values farther in that direction until performance decreased, or if performance improved before declining for values in a particular direction, we chose the parameter value that produced the optimal validation performance. We then trained the model using all of the optimal parameter values together. After comparing the performance of each of the model configurations, including the models for which one variable was changed at a time as well as the model combining all of the optimal parameter values, we selected the best combination of parameter values to be that with the best performance. We explored 3-5 values for each parameter, resulting in approximately 25-35 configurations per model.

All parameter tuning was done using the portion of the data reserved for training and validation. Each configuration was trained on the training portion of the five train/validation folds and evaluated on the held-out validation fold. The average performance across these train/validation splits was used to determine the best parameter configuration while accounting for sampling noise. After all parameters were tuned, final performance was reported on the reserved test dataset which was not seen by the model during training nor parameter tuning.

### 5.3 Baseline Methods

We compare our method to multiple baselines, which were used to generate embeddings from multi-modal data in the first part of the pipeline. These embeddings were subsequently input to the prediction models mentioned above. The baselines include the original data, deep canonical correlation analysis (DCCA) [27], generalized canonical correlation analysis (GCCA) [28], multimodal variational autoencoder (MVAE) [13], cross-linked unified embedding (CLUE) [12], and IntegrAO [15].

Original data consists of concatenating the original features across all modalities and treating the resulting vector as the input to the prediction model. The canonical correlation analysis (CCA)-based methods serve as baseline dimensionality reduction methods that produce unified embeddings incorporating information across multiple data sources. Generally, CCA aims to find linear projections of two data sources with maximal correlation, and thus can learn relevant inter-modality relationships. DCCA extends CCA to learn nonlinear relationships but is limited to two modalities at a time. To use DCCA, we learn all pairwise projections and then take the average across all learned projections for each modality before concatenating the averaged projections across modalities. GCCA extends CCA to handle more than two data sources at a time but is still limited to linear projections. For the original data, DCCA, and GCCA, which do not handle missing modalities, we subset the samples to those that have all modalities.

On the other hand, MVAE and CLUE are autoencoder-based methods that can handle missing modalities. MVAE does this using a product-of-experts approach to define a joint embedding. CLUE, on the other hand, does not learn a joint embedding but instead uses self- and cross-encoders to learn modality-specific representations while being able to impute missing modalities. IntegrAO is a recently published graph-based method that constructs omics-specific patient graphs and combines them utilizing graph neural networks to produce unified patient embeddings. It utilizes patient similarity to propagate information to subjects with missing modalities from those that are similar. Since these methods can handle missing modalities, they were trained on all samples. To keep the testing data consistent across all methods for fair comparison, test results for all methods were reported for only the samples with all modalities. However, we note that it is possible for MVAE, CLUE, IntegrAO, and JASMINE to perform inference on samples with missing modalities. A summary of these methods and their features is presented in Table 1.

### 5.4 Evaluation Metrics

#### Survival prediction performance metrics

For survival prediction, we used the concordance index for right-censored data based on inverse probability of censoring weights (C-index IPCW) and cumulative dynamic area under the receiver operating characteristic curve (CD AUC). C-index IPCW measures the how well the method predicts the ordering of the survival times while handling a large proportion of censoring (69.4% of the samples in TCGA are censored) by using the inverse probability of censoring weights. It does not depend on the distribution of censoring times in the test data and thus can produce unbiased estimates [98]. CD AUC is analogous to the AUROC in that it converts the inference to a binary classification problem by predicting survival at a given cutoff time and assesses the sensitivity and specificity of the predictions across a range of time thresholds.

#### Classification performance metrics

For the classification tasks, we evaluated the performance using balanced accuracy, AUROC and average precision score. Balanced accuracy measures the correctness of the predicted classes while weighting each sample by the inverse prevalence of its true class. For AUROC, we used the macro averaged one-versus-rest AUROC across all classes. Average precision score (AP) is analogous to the AUROC in that it summarizes the precision-recall curve by taking the weighted mean of precisions achieved across a range of predicted probabiity thresholds. Weighting is determined by the increase in recall from the previous threshold [98]. For the multiclass scenario, the macro-average across classes is utilized.

#### Clustering performance metrics

We applied K-means clustering to the embeddings over a range of cluster numbers. In our experiments, we varied the number of clusters from 3 to 8. In TCGA, we tested for differential survival among clusters of TCGA subjects within five different cancer types. For this, we used the multivariate log-rank test, which tests for differences in survival curves among multiple clusters. We reported − log 2(*p*) of this test, where *p* is the p-value

#### Regression performance metrics

For regression tasks, we used root mean squared error (RMSE), the coefficient of determination (*R*^2^-score), and the Pearson correlation between the predicted and true values (r-pred). RMSE takes the square root of the mean of squared differences between the true and predicted values across samples. The *R*^2^-score, as computed using the r2 score function in scikit-learn [98], measures the proportion of variation in the data that is explained by the model, i.e., what is left after subtracting the proportion of variation in the data captured by the error in predictions:

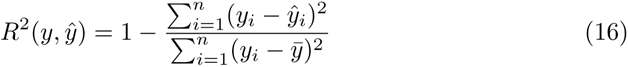

where *y* is the true outcome, *ŷ* is the predicted outcome, and *y̅* is the expected value of *y*. This measures how well future samples are likely to be predicted by the model, where 1.0 is the best possible score, and the model that always predicts *y̅* has an *R*^2^ score of 0 [98].

### 5.5 Feature Importance Analysis

JASMINE itself is a self-supervised representation learning method, and thus the embeddings it generates are not specialized for a specific task. Furthermore, employment of JASMINE for a particular prediction problem involves a two-step pipeline: first generating the embeddings, and then inputting the representations to a down-stream prediction model. To identify the features in each modality driving a particular prediction, we also approach feature interpretation in a two-step manner. This involves first using the downstream prediction model to identify which embedding features are most important for the prediction task. Then, once the top embedding features are identified, we interpret these features by identifying which original interpretable variables (e.g. genes, proteins, miRNAs) from each modality contribute most to those embedding features.

In our experiments, we focused on the cancer survival prediction task. The down-stream predictive model for survival prediction was the Coxnet regression model. We first selected the top predictive embedding feature determined by this model by identifying the feature corresponding to the regression coefficient with the largest absolute value.

Each element of the embedding is the result of complex functions of the original inputs and is thus not directly interpretable. To elucidate its potential biological underpinnings, we identified the original interpretable features with the greatest influence on the embedding feature of interest. We determined this by applying a perturbation via shuffling to the original features and measuring the impact on the embedding feature of interest. This idea is based on permutation feature importance [109, 110], which identifies a feature as important if shuffling its values meaningfully increases prediction error. Since JASMINE is a self-supervised representation learning method, we modified this approach to measure the change in the embedding feature rather than prediction error. This involved two main steps: (1) identifying groups of correlated features within each modality, and for each group, (2) permuting the values of all of its constituent features and measuring the impact on the embedding feature of interest. The features within each modality that produced the greatest change in the embedding feature were interpreted to be the most important contributors to that embedding feature.

Because many of the input features are correlated, we first identified groups of correlated features within each modality before applying the permutation to these feature groups rather than individual features. This served two main purposes: (1) it reduced the computational burden of testing tens of thousands of individual features to testing thousands of feature groups, and (2) it helped avoid the case in which correlated features capturing similar information reduce the impact of perturbing a single important feature, which could result in overlooking truly important features.

To generate groups of correlated features, we computed the correlation matrices between the features in each modality separately. Then, we converted these correlation matrices to distance matrices by subtracting the absolute values of their elements from 1. We then applied agglomerative clustering to the distance matrices to identify clusters of correlated features within each modality [111–113].

For the perturbation step, we applied the shuffling at the cluster-level by permuting the values of all features within a given cluster before evaluating the impact on the embedding feature of interest. This was calculated using the correlation across samples between the value of the generated embedding feature before and after shuffling the input features within the cluster of interest. We expect that if an input feature is more relevant to the embedding feature, then permuting its values will have a greater impact on the value of the embedding feature, resulting in a lower correlation. We used correlation rather than a measure of magnitude of change to account for differences in scales among the features: a larger scale feature may produce a greater shift in the embedding feature than a smaller scale feature of equal importance.

Overall, this procedure resulted in one group of features for each modality that was designated as most important for a given embedding feature. We then performed pathway and disease enrichment analyses on these features to interpret the underlying biology captured by the embedding. For the top mRNAseq, Methylation, and RPPA gene clusters, we used PANTHER [30] for GO Enrichment Analysis from the Gene Ontology Consortium [31, 32] to determine the top Reactome [33] pathways represented by these genes. The miRNAseq features were miRNA molecule IDs rather than genes; thus, for the top miRNA cluster, we used the miRNA Enrichment Analysis and Annotation Tool (miEAA) [34] for over-representation analysis to identify the top associated diseases from the Human microRNA Disease Database (HMDD) [35]. Additionally, we analyzed overlapping information among the modalities to evaluate whether the embedding captured biologically meaningful inter-modal feature correspondence. For mRNAseq, Methylation, and RPPA, we identified the genes within the top feature clusters that were shared by at least two modalities. For miR-NAseq, we identified the target genes for each miRNA molecule in the selected feature cluster that overlapped with at least two of the other modalities. The target genes and their target prediction scores for each miRNA were obtained from miRDB [70, 71]. Target prediction score ranges between 50-100, with higher score indicating more confidence in the prediction. We then assessed the overlapping genes for associations with a prediction task of interest.

### 5.6 Data processing

The data were split into train, test, and validation sets. We used multiple train/validation splits for tuning hyperparameters and assessing performance variability. For our experiments, there were five train/validation splits, each resulting in one trained model, and a single test dataset. We report the means and standard deviations of the performance metrics on the test data from the five models. For more details on the dataset-specific splits and hyperparameter settings, see Sections 5.6.2 and 5.6.3.

Our experimental pipeline consists of two main steps: (1) generate embeddings using an unsupervised representation learning method and (2) input the embeddings to a prediction model of choice for a given inference problem. To generate embeddings for part (1) of our pipeline, we first trained the models on each training dataset and then used them to generate embeddings for all data samples. For part (2) of the pipeline, we used the embeddings of the training data to train a predictive model on a specific task (e.g. survival prediction, classification, regression). Performance of the predictive model on the embeddings of the validation data were used to tune the hyperparameters of the models. Final results were reported on the embeddings of the test data.

#### 5.6.1 Simulated data

To compare JASMINE and the baseline methods under controlled data scenarios, we applied them to simulated data. These data were simulated as coming from three different classes, each consisting of 3,000 samples for a total sample size of 9,000. For each sample, informative features which distinguished class labels were generated from a multivariate Normal distribution. We simulated six modalities with 300 informative features per modality. Some experiments involved fewer than six modalities; for these, we simply ignored the extra modalities. Our simulated data incorporated both modality-shared and specific informative features. Modality-shared features had a correlation of 0.3 with the corresponding features from all other modalities, whereas modality-specific features had a correlation of 0.

To generate the data, we first sampled class centroids from a multivariate Normal distribution with zero mean and inter-class variances of 0.1. These centroids represented all informative features from all modalities, for a total of 300 × 6 = 1, 800 informative features. The correlations among corresponding features of different modalities were set based on whether the features were modality-specific (correlation of 0) or modality-shared (correlation of 0.3). Once the class centroids were generated, we produced individual samples by taking 3,000 random draws from a multivariate Normal distribution with the class centroid as the mean and intra-class variance of 0.3. Again, the correlations among features were set based on whether the features were modality-specific or modality-shared. Once the total feature vectors were generated, we split them into feature sets representing different modalities. For each modality, we generated 1,800 uninformative features from a standard uniform distribution to represent noise in the data. Finally, we applied a nonlinear mapping by squaring the data so that they were not linearly separable. We also generated an additional dataset using the same settings but reducing the number of samples per class to 1,500 for a total sample size of 4,500 for experiments at small sample size.

Our simulation experiments involved altering the modality missingness patterns and the number of modalities. To do this, we generated random masks to indicate which modalities are missing for each sample. Masks were boolean, with True indicating that a modality is available and False indicating that it is missing. We generated masks for three types of experiments: (1) balanced missingness at various modality missingness rates, (2) unbalanced missingness based on class, and (3) varying the number of modalities.

For balanced missingness at various rates, every modality had the same probability of being missing for all samples, regardless of their true labels. The total number of modalities was four. For each simulation setting, we varied proportion of samples with one, two, three, and four modalities available. The specific proportions for each modality missingness rate are the following:

- For the missingness rate of 0, 100% of the samples had four modalities.
- For the missingness rate of 0.38, 12% had one modality, 38% had two, 38% had three, and 12% had four.
- For the missingness rate of 0.53, 50% had one modality, 20% had two, 20% had three, and 10% had four.
- For the missingness rate of 0.66, 80% had one modality, 8% had two, 8% had three, and 4% had four.
- For the missingness rate of 0.74, 97% had one modality, 1% had two, 1% had three, and 1% had four.

For each setting, we randomly split samples into four groups designating whether they have one, two, three, or four modalities available. Then, for each sample we generated a boolean mask of length equal to the number of modalities, with the number of ‘True’ values corresponding to the number of modalities available for that sample. Finally, we shuffled the mask for each sample so the missing modalities were randomly chosen, resulting in balanced missingness patterns.

For unbalanced missingness based on class, we simulated the case when one modality was more likely to be missing, and this modality was chosen based on the sample’s true class. Let *p* denote the missingness probability of a modality when it is ‘more likely’ to be missing, and let *q* be the missingness probability otherwise. The per-modality missingness rates were chosen as follows: samples in class 1 were missing modality 1 with probability *p* but missing the other modalities with probability *q*. Samples in class 2 were missing modality 2 with probability *p* but missing the others with probability *q*. Samples in class 3 were missing modality 3 with probability *p* but missing the remaining modalities with probability *q*. The total number of modalities was four. We varied *p* and *q* to produce a range of overall missingness rates: 0.18, 0.28, 0.38, 0.48, 0.58. The specific values of *p* and *q* for each of these settings are the following:

- For the overall missingness rate of 0.18, *p* = 0.6 and *q* = 0.1.
- For the overall missingness rate of 0.28, *p* = 0.7 and *q* = 0.2.
- For the overall missingness rate of 0.38, *p* = 0.8 and *q* = 0.3.
- For the overall missingness rate of 0.48, *p* = 0.9 and *q* = 0.4.
- For the overall missingness rate of 0.58, *p* = 0.99 and *q* = 0.5.

Finally, for the experiments varying the number of modalities, all samples were simulated to have modality 1 available. For each following modality, the probability of missing that modality was 0.95. We generated masks independently for each modality, each with the same missingness probability across all samples. The total number of modalities ranged from two to six.

#### 5.6.2 TCGA

We applied JASMINE and the baseline methods to multi-omics data from The Cancer Genome Atlas (TCGA) [29], consisting of mRNA-seq, methylation, miRNA-seq, and functional proteomics reverse phase protein array (RPPA) modalities.

The mRNA-seq data were obtained using the Affymetrix HG U133, Affymetrix Exon Array and Agilent gene expression platform. Each feature corresponds to the transcript abundance calculated from microarray intensities for a given gene (e.g. CA5BP, ERP27, TP53).

The methylation data represent the normalized methylation *β* values calculated from methylated and unmethylated probe intensities, quantifying methylation levels at each gene. Thus, each feature corresponds to a gene (e.g. BRCA1, FOXA1, GRP). Genes on chromosomes X or Y, with gene symbol NA, and for which more than 5% of the *β* values were NA were filtered out.

The miRNAseq data were RPKM log2 transformed gene-level counts calculated from microarray intensities. Each feature represents a gene (e.g. hsa-mir-1247, hsamir-216a, hsa-mir-770). All records with NA values were removed.

The RPPA data represent protein expression levels measured via antibody probe and calculated from spot intensities. Each feature corresponds to a gene/protein (e.g. CLDN7|Claudin-7, ERBB3|HER3, CTNNB2|beta-Catenin). Features with NA values for any sample were removed.

All features were standardized to have zero mean and standard deviation of 1 before being input to the models. This ensured that the features were on a similar scale within and across modalities.

This dataset consists of 10,472 total samples, some of which are missing one or more modalities in a blockwise fashion, i.e., all features for a modality are either observed or not observed. We kept the features that were measured for all subjects with the corresponding modality. To test our methods’ ability to handle a large number of features, we did not perform any prior feature selection or dimensionality reduction. The summary of sample sizes, dimensionality, and labels are available in Table 2.

For TCGA, we split the data samples into six folds, five of which were used for 5-fold cross-validation. The sixth split was used as a held-out test dataset, on which our final results are reported. For the methods that do not handle missing data (original data, DCCA, GCCA), we subset the data to samples with all modalities available (6,054 samples). For the remaining baseline methods (MVAE, CLUE, IntegrAO) and JASMINE, we use all samples for training.

For cancer stage classification, we used pathologic stage as the label, which is based on clinical staging and histopathologic examination of surgical specimens to determine the extent of cancer spread [114]. There are four main cancer stages, I, II, III and IV, each additionally comprising sub-stages. We removed samples with stages that did not fall into any of these categories due to very little representation of these categories in the data. We aggregated the sub-stages into the four main stages as the labels for four-class classification.

For training, we used the RMSprop optimizer from PyTorch, learning rate of 0.002 and dropout of 0.2. The model architecture consisted of the following parameter settings: encoder depth of 2, encoder hidden dimension of 512, decoder depth of 1, decoder hidden dimension of 256. The dimension of the learned representations was set to 64 (this setting was used for all baseline methods, except for original data). The best settings for the weights of the loss terms included in our method, which were deter-mined based on performance on the validation data, were found to be *λ_self_* = 1.0, *λ_cross_* = 1.0, *λ_cosine_* = 3.0, *λ_joint_* = 1.0, *λ_MCL_* = 1.0, *λ_SCL_*= 0.5, and *λ_orthog_*= 0.5.

#### 5.6.3 ADNI

We also applied JASMINE to multi-omics data from the Alzheimer’s Disease Neuroimaging Initiative (ADNI) [96, 97]. Data used in the preparation of this article were obtained from the Alzheimer’s Disease Neuroimaging Initiative (ADNI) database (adni.loni.usc.edu). The ADNI was launched in 2003 as a public-private partnership, led by Principal Investigator Michael W. Weiner, MD. The original goal of ADNI was to test whether serial magnetic resonance imaging (MRI), positron emission tomography (PET), other biological markers, and clinical and neuropsychological assessment can be combined to measure the progression of mild cognitive impairment (MCI) and early Alzheimer’s disease (AD). The current goals include validating biomarkers for clinical trials, improving the generalizability of ADNI data by increasing diversity in the participant cohort, and to provide data concerning the diagnosis and progression of Alzheimer’s disease to the scientific community. For up-to-date information, see adni.loni.usc.edu.

From ADNI, we utilized gene expression (GEX), genetic (GEN), metabolomics (MET), and proteomics (PROT) modalities. The GEX data were microarray gene expression profile data calculated from microarray intensities for a given transcript. Expression values were preprocessed via the RMA (Robust Multi-chip Average) normalization method. Each feature corresponds to a transcript, given by the Affymetrix probe ID (e.g. 11715373_a_at, 11723370_s_at, 11729407_a_at). Although the ADNI GEX data are mRNA-seq measurements, they are not equivalent to the mRNA-seq data in TCGA, as these represent probe-level rather than gene-level expression, and they were preprocessed using different normalization methods. The GEN data are allele counts for 935 SNPs derived from key AD GWAS studies [115, 116]. These counts were extracted via PLINK.

The MET data were flow injection analysis (FIA) and ultra performance liquid chromatography (UPLC) data from which represent serum metabolite concentrations. The FIA data were measured for the Glycerophospholipid, Sphingolipid, and Acylcarnitine analyte classes; the UPLC data were measured for the Amino Acid and Biogenic Amine analyte classes. These concentrations were calculated following peak integration and calibration of the mass spectrometry data. Each feature represents a metabolite (e.g. C0, C14.1, C16 for FIA and Ala, Arg, Asn for UPLC).

The PROT data represent median normalized CSF protein levels measured via SomaScan 7K assay and calculated from probe intensities. Each feature represents an analyte ID (e.g. X10589.7, X16916.19, X3336.50), which is convertible to its corre-sponding gene or protein ID. Feature filtering performed by ADNI included removing aptamers based on thresholding the scale factor, the coefficient of variation, and call rate as well as removing outliers, non-human analytes, and analytes without protein targets. For both the MET and PROT data, features that were missing more than 5% of the samples were removed; for the remaining features, missing values were imputed using the mean of that feature across all samples.

All features were standardized to have zero mean and standard deviation of 1 before being input to the models. This ensured that the features were on a similar scale within and across modalities.

The dataset comprises a total of 1,665 samples, including several that have at least one missing modality. As with the TCGA data, missingness is blockwise such that either all features or no features are missing for a given modality. For gene expression and genetic data, all features were measured for all samples. For metabolomics and proteomics data, we dropped features that were measured for fewer than 95% samples. For the remaining features, we filled missing values with the sample means. For samples with repeated proteomics measurements, we utilized the first measurement available. Baseline diagnoses and cognitive scores were extracted for each patient. The majority of metabolomics and proteomics samples were baseline measurements. Many gene expression measurements were not taken at baseline, but we included these measurements as a proxy for baseline gene expression. The sample sizes, features, modalities, and labels are summarized in Table 2.

For ADNI, due to the smaller sample size, we split the data first into 70% training and 30% testing. Then, within the training set, we generated five random 70%/30% training/validation splits. As with TCGA, for the methods that do not handle missing data (original data, DCCA, GCCA), we subset the data to samples with all modalities available (230 samples). For the remaining baseline methods (MVAE, CLUE, IntegrAO) and our proposed method (JASMINE), we use all samples for training. For fair comparison, all methods are tested only on samples with all modalities.

We applied JASMINE to diagnosis classification and MMSE (Mini-Mental State Examination) prediction. The data contained three main diagnoses: CN (cognitively normal), MCI (mild cognitive impairment), and AD (Alzheimer’s dementia). Due to small sample size and class imbalance resulting in relatively few AD samples, we converted the task to a binary classification problem, with CN and MCI/AD as the two classes. MMSE is a cognitive exam consisting of 30 questions that evaluate attention and orientation, memory, registration, recall, calculation, language and ability to draw a complex polygon [117]. Lower scores indicate worse cognitive state, and a score less than 24 has traditionally been used as a cutoff indicating evidence of dementia. As the score varies from 0 to 30, we posed this as a regression task.

For training, we used the RMSprop optimizer from PyTorch, learning rate of 0.002 and dropout of 0.2. The model architecture consisted of the following parameter settings: encoder depth of 2, encoder hidden dimension of 512, decoder depth of 1, decoder hidden dimension of 256. The dimension of the learned representations was set to 64 (this setting was used for all baseline methods, except for original data). The best settings for the weights of the loss terms included in our method, which were determined based on performance on the validation data, were found to be *λ_self_* = 1.0, *λ_cross_* = 1.0, *λ_cosine_* = 3.0, *λ_joint_* = 1.0, *λ_MCL_* = 2.0, *λ_SCL_*= 0.5, and *λ_orthog_*= 0.5.

## 6 Declarations

### Ethics approval and consent to participate

Not applicable

### Consent for publication

Not applicable

### Availability of data and materials

The TCGA multi-omics data are available in the Broad GDAC Firehose repository, https://gdac.broadinstitute.org/ [118]. Clincal data were files ending in ‘.clin.merged.picked.txt’ from the Clinical_Pick_Tier1. mRNA-seq data were files ending in ‘.uncv2.mRNAseq_RSEM_all.txt’ from the mRNAseq_Preprocess. Methylation data were files ending in ‘.meth.by_mean.data.txt’ from the Methylation_Preprocess. miRNA-seq data were files ending in ‘.miRseq_RPKM log2.txt’ from miRseq_Preprocess. RPPA data were from files ending with ‘.rppa.txt’ from RPPA_AnnotateWithGene.

The ADNI data are available in the Laboratory of Neuroimaging (LONI) Image & Data Archive (IDA), https://ida.loni.usc.edu/login.jsp [119], but restrictions apply to the availability of these data, which were used under license for the current study, and so are not publicly available. Baseline diagnoses and cognitive scores were taken from the ADNIMERGE data file for ADNI1/GO/2/3. Gene expression data were taken from the Microarray Gene Expression Profile Data. Genetic data were extracted for 935 SNPs derived from key AD GWAS studies [115, 116]. Metabolomics data were from the ADMC Duke Biocrates p180 Kit Flow injection analysis [ADNIGO,2] and ADMC Duke p180 Ultra Performance Liquid Chromatography [ADNIGO,2]. Proteomics data were from CruchagaLab CSF SOMAscan7k Protein matrix postQC [ADNI1,GO,2].

The code for JASMINE is available under the MIT license on GitHub with the URL https://github.com/jballard28/JASMINE. The software is platform independent and requires Python 3.8 or higher.

### Competing interests

The authors declare that they have no competing interests.

### Funding

Q.L. discloses support for the research of this work from NIH/NIA [grant numbers R01-AG071174 and RF1-AG063481], and NIH/NCI [grant number U01 CA274576]. L.S. discloses support for the research of this work from NIH/NIA [grant numbers U01-AG068057, U01 AG066833, R01-AG071470 and U19-AG074879].

### Authors’ contributions

J.L.B., Z.D., and Q.L. conceived the project. J.L.B. and Z.D. developed the methods and implemented the baseline methods. Z.D. conducted initial method implementation and experiments. J.L.B. executed all the analyses, including data preparation and preprocessing, method refinement and implementation, and numerical experiments. J.L.B., Q.L., and L.S. contributed to the design of the numerical experiments and the interpretation of the results. J.L.B. generated the figures and wrote the manuscript with input from all authors. All authors contributed to editing and revising the manuscript. Q.L. and L.S. jointly supervised the project.

## Acknowledgments

Data collection and sharing for the Alzheimer’s Disease Neuroimaging Initiative (ADNI) is funded by the National Institute on Aging (National Institutes of Health Grant U19AG024904). The grantee organization is the Northern California Institute for Research and Education. In the past, ADNI has also received funding from the National Institute of Biomedical Imaging and Bioengineering, the Canadian Institutes of Health Research, and private sector contributions through the Foundation for the National Institutes of Health (FNIH) including generous contributions from the following: AbbVie, Alzheimer’s Association; Alzheimer’s Drug Discovery Foundation; Araclon Biotech; BioClinica, Inc.; Biogen; BristolMyers Squibb Company; CereSpir, Inc.; Cogstate; Eisai Inc.; Elan Pharmaceuticals, Inc.; Eli Lilly and Company; EuroImmun; F. Hoffmann-La Roche Ltd and its affiliated company Genentech, Inc.; Fujirebio; GE Healthcare; IXICO Ltd.; Janssen Alzheimer Immunotherapy Research & Development, LLC.; Johnson & Johnson Pharmaceutical Research & Development LLC.; Lumosity; Lundbeck; Merck & Co., Inc.; Meso Scale Diagnostics, LLC.; NeuroRx Research; Neurotrack Technologies; Novartis Pharmaceuticals Corporation; Pfizer Inc.; Piramal Imaging; Servier; Takeda Pharmaceutical Company; and Transition Therapeutics.

Data collection and sharing for this project was funded by the Alzheimer’s Disease Metabolomics Consortium (National Institute on Aging R01AG046171, RF1AG051550 and 3U01AG024904-09S4).

## Notes

https://github.com/jballard28/JASMINE

### Competing Interest Statement

The authors have declared no competing interest.

### Summary of Updates

Simulation experiments updated with more realistic missingness patterns and setting names updated to clarify and better represent overall modality missingness rate. More settings for the unbalanced missingness experiments were added. Figure 2 updated with the new simulation results.

